# T-cell receptor *αβ* chain pairing is associated with CD4^+^ and CD8^+^ lineage specification

**DOI:** 10.1101/293852

**Authors:** Jason A. Carter, Jonathan B. Preall, Kristina Grigaityte, Stephen J. Goldfless, Adrian W. Briggs, Francois Vigneault, Gurinder S. Atwal

**Affiliations:** Department of Applied Mathematics and Statistics, Stony Brook University, Stony Brook, NY 11794; Cold Spring Harbor Laboratory, Cold Spring Harbor, NY 11724; Watson School of Biological Sciences, Cold Spring Harbor Laboratory, Cold Spring Harbor, NY 11724; Juno Therapeutics, Seattle, WA 98109

## Abstract

While a highly diverse T-cell receptor (TCR) repertoire is the hallmark of a healthy adaptive immune system, relatively little is understood about how the CD4^+^ and CD8^+^ TCR repertoires differ from one another. We here utilize high-throughput single T-cell sequencing to obtain approximately 100,000 TCR *αβ* chain pairs from human subjects, stratified into CD4^+^ and CD8^+^ lineages. We reveal that substantial information about T-cell lineage is encoded by V*αβ* gene pairs and, to a lesser extent, by several other TCR features such as CDR3 length and charge. We further find that the strength of association between the *β* chain and T-cell lineage is surprisingly weak, similar in strength to that of the *α* chain. Using machine learning classifiers to predict T-cell lineage from TCR features, we demonstrate that *αβ* chain pairs are significantly more informative than individual chains alone. These findings provide unprecedented insight into the CD4^+^ and CD8^+^ TCR repertoires and highlight the importance of *αβ* chain pairing in TCR function and specificity.

## 1. Introduction

During thymic positive selection, bipotent T-cell precursors differentiate into either CD4^+^ helper T-cell or CD8^+^ cytotoxic T-cell lineage. While this process is contingent upon the interaction of the heterodimeric *αβ* T-cell receptor (TCR) with either MHC class II or I, respectively, relatively little is currently known about the TCR features mediating this interaction^1–3^. One possible explanation posits the existence of germline-encoded sequences that have been hard-wired into the Variable (V) region’s CDR1 and CDR2 loops^4–13^. Recent support for such germline-bias includes the finding that expression levels of specific TCR V-regions are correlated with MHC polymorphisms ^14^. However, the role of the entire *αβ* chain sequence in specifying CD4^+^ and CD8^+^ repertoires has remained unknown.

While previous methods for paired *αβ* TCR sequencing have been developed ^15–21^, only recently have technological advances enabled high-throughput capture of paired *αβ* TCR sequences^22–25^. As both *α* and *β* chains have been implicated to play important roles in

TCR binding of the peptide-MHC (pMHC) complex, it follows that such single-cell sequencing methods may reveal differences in the paired TCR repertoires between each T-cell lineage^26–32^. Thus, in order to better understand the factors that influence T-cell differentiation, we addressed how the paired *αβ* TCR repertoires differ between the CD4^+^ and CD8^+^ T-cell populations.

## 2. Results

### Overlap between the CD4^+^ and CD8^+^ repertoires

We previously employed a novel high-throughput, single-cell sequencing method to capture TCR pairs obtained from the peripheral blood of 5 healthy individuals^24,33^. In this study, we utilized another single-cell microfluidic platform (10x Genomics) ^25^ to add to this database and create the largest database of paired CD4^+^ and CD8^+^ TCR sequences to date (Sup. Figs. 1 and 2). Using this dataset comprised of nearly 100,000 paired *αβ* TCR sequences, we first assessed the CD4^+^ and CD8^+^ TCR repertoire overlap.

Considering the unique set of TCR clonotypes (V*αβ* and amino acid CDR3*αβ*) across all individuals, we found that the paired CD4^+^ and CD8^+^ repertoires were largely disjoint from one another. Splitting the paired repertoire into the constituent *α* and *β* populations resulted in considerably higher overlap between the two lineages (Fig. 1A-C). Next quantifying the overlap between the CD4^+^ and CD8^+^ TCR repertoires within each individual, we observed greater similarity between the CD4^+^ and CD8^+^ single chain repertoires than between the paired *αβ* repertoires (Fig. 1D). Previous findings have suggested that TCRs shared between individuals may have shorter CDR3*β* sequences^34^ and may be closer to germline recombination sequences than clonotpyes found only in a single individual ^35–37^. Accordingly, TCR sequences shared between the CD4^+^ and CD8^+^ lineages were, on average, shorter than those found only in one of the two lineages with respect to the *α* (*p*=1.4×10^−5^), *β* (*p*=6.3×10^−8^) and *αβ* (*p*=9.3×10^−6^ by Mann-Whitney U test) repertoires(Fig. 1E and Sup. Fig. 3).

**Figure 1:**
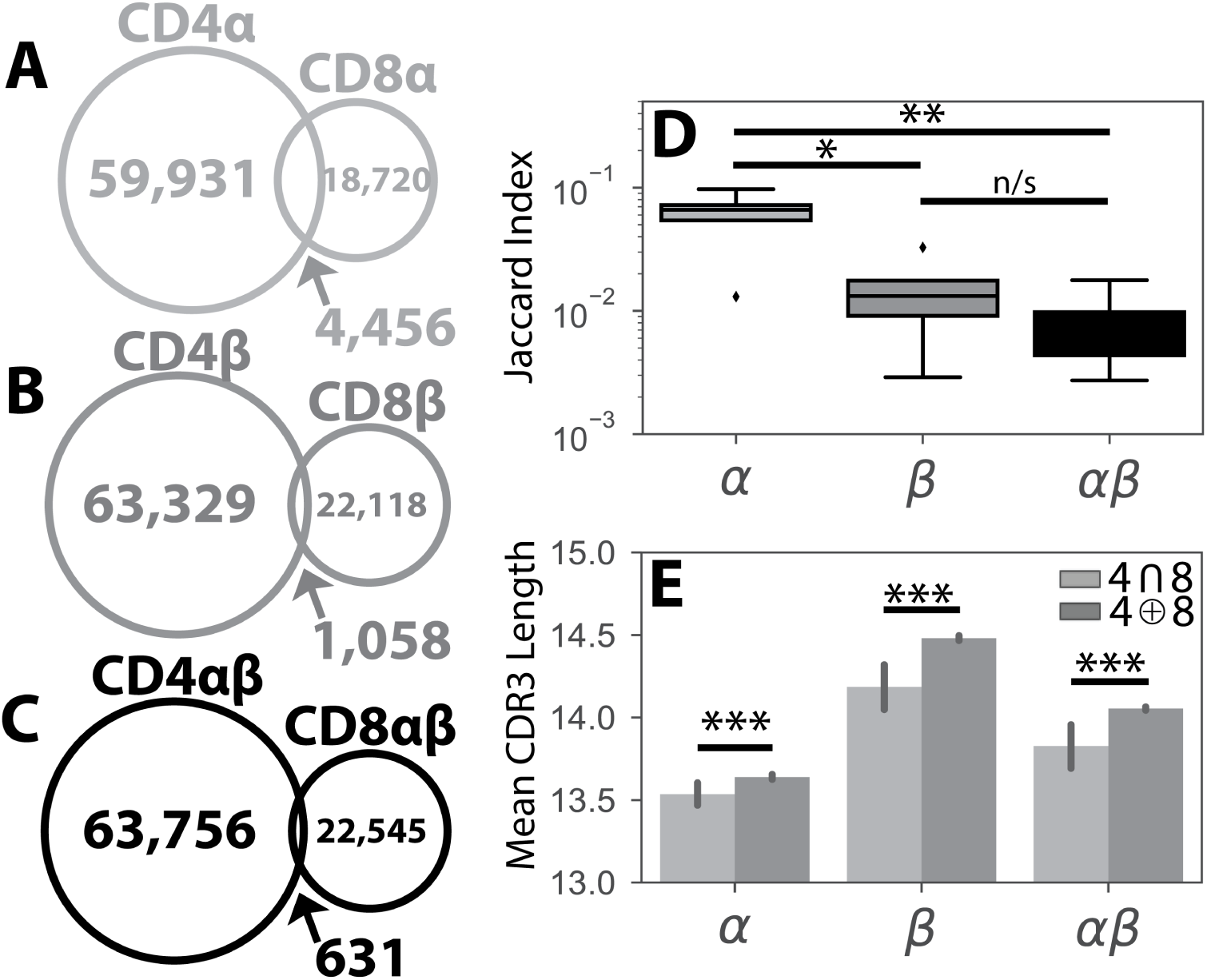
Overlap between CD4^+^ and CD8^+^ TCR repertoires. Population unique TCRs, defined as V*αβ*-CDR3*αβ* amino acid sequence clonotypes, were calculated by combining sequences from all 6 individuals separately for the CD4^+^ (n=64,387) and CD8^+^ (n=23,176) subsets. The overlap between the global CD4^+^ and CD8^+^ repertoires were then calculated for the **(A)** *α*, **(B)** *β*, and **(C)** paired *αβ* repertoires. **(D)** The Jaccard Index, a measure of similarity between 2 sets, was calculated pairwise for each individual’s CD4^+^ and CD8^+^ *α*, *β* and paired *αβ* repertoires. Significance between samples was assessed using a Student’s t-test. **(E)** Bar plots show mean CDR3 lengths for *α*, *β*, and *αβ* TCR sequences found exclusively in (dark gray, ⊕) or shared between (light gray, ∩) the CD4^+^ and CD8^+^ lineages. Error bars represent bootstrapped 99% confidence intervals for the mean. Shared sequences were significantly shorter than those sequences found in both repertoires by Mann-Whitney U Test. For all panels, n/s-not significant, ^*^*p<*0.05, ^**^*p<*0.01, and ^***^*p<*0.001.

The decreased CD4^+^ and CD8^+^ repertoire overlap for *αβ* pairs relative to either single chain repertoire may reflect an increased specificity of *αβ* pairs for a given MHC class. As this explanation would be biologically consistent with previous structural findings implicating both chains in determining TCR-pMHC binding^26–32^, we further explored the extent to which *αβ* pairs could be used to provide additional information on T-cell lineage as opposed to the either chain alone.

### Association of VJ germline segment usage with CD4^+^-CD8^+^ status

Significant biases in V and J germline segment use between the single-chain CD4^+^ and CD8^+^ repertoires have been identified previously^38–40^. To further explore this, we calculated the frequency with which all V*α* and V*β* regions were used by each individual (Fig. 2A-B). While variations in the usage statistics exist between individuals, our results are in general agreement with previous estimates (Sup Figs. 4-7) ^41,42^. The association between each V region and T-cell lineage was quantified by calculating the odds ratio^38^, revealing only weak associations between the usage of a particular V*α* or V*β* segment and T-cell lineage (Fig. 2C-D). Weaker associations between T-cell lineage and single chain J*α* and J*β* usage were also present (Sup. Fig. 8A-D). Interestingly, these associations for both V- and J-regions are significantly weaker than previously reported^38^.

**Figure 2:**
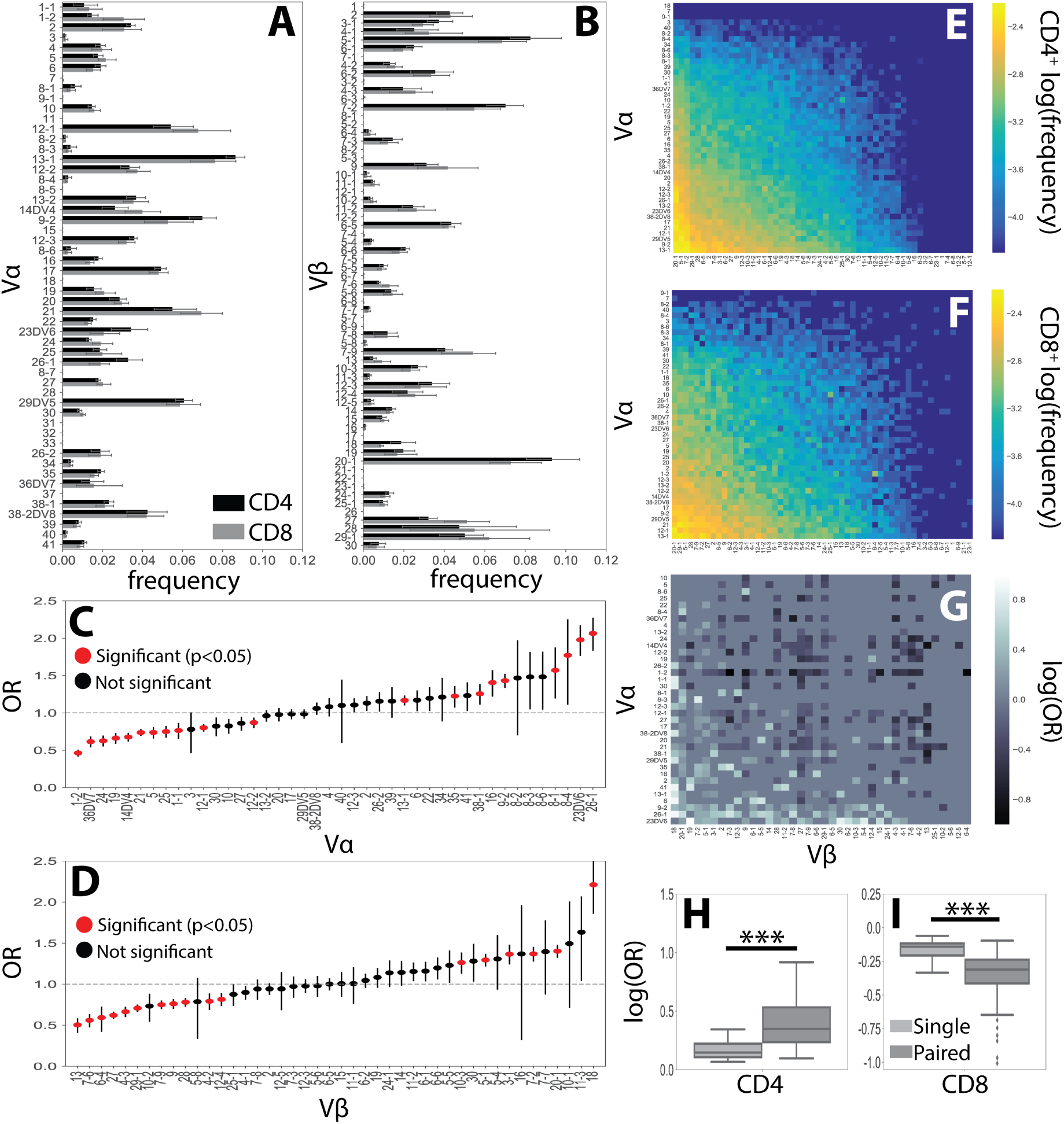
V germline region usage in the *α*, *β* **and** *αβ* **repertoires**. **(A)** V*α* and **(B)** V*β* single-chain germline region usage frequencies were calculated for each individual’s CD4^+^ and CD8^+^ T-cell repertoires. Error bars represent the standard deviation across individuals. **(C)** The CD4^+^ (n=63,718) and CD8^+^ (n=22,534) TCR repertoires were then pooled across individuals and the CD4^+^:CD8^+^ odds ratio (OR) was calculated for each V*α* and **(D)** V*β* germline region. An OR*>* 1 represents a CD4^+^ bias, while an OR*<* 1 represents a CD8^+^ bias with error bars representing the 95% confidence interval. The mean is represented by a red or black dot, with red representing statistical significance at *p<*0.05 by Fisher’s exact test after applying Bonferroni correction. **(E)** Paired V*αβ* usage frequencies across all individuals for CD4^+^ and **(F)** CD8^+^ TCR repertoires. **(G)** Significant (*q<*0.05 by Fisher’s exact test) log odds ratios reveals strong CD4^+^:CD8^+^ biases for 355 V*αβ* pairs. **(H)** Boxplots were calculated for the set of all significant odds ratios associated with single chains (V*α* or V*β*) and compared with those associated with *αβ* pairs. Paired associations for both CD4^+^ and **(I)** CD8^+^ status were significantly stronger (\*\*\**p<* 0.001 by Mann-Whitney U test) than those associated with a single chain alone.

The role of paired germline segment usage in biasing T-cell differentiation was examined by comparing the V*αβ* and J*αβ* paired distributions for both T-cell populations (Fig. 2E-F and Sup Fig. 8E-F). The CD4^+^:CD8^+^ odds ratio was then calculated for each germline pair (Sup. Figs. 9-11). Our results reveal 352 V*αβ* and 70 J*αβ* pairs associated with a significant (*q<*0.05) lineage specification bias (Fig. 2G and Sup Fig. 8G). Interestingly, the strength of association with T-cell lineage was significantly stronger for V*αβ* pairs than for J*αβ* pairs, likely reflecting the contribution of the CDR1 and CDR2 loops present in each V region to MHC binding^43^.

We further note the association between paired V*αβ* and cell lineage was significantly stronger (CD4^+^: *p*=2.1×10^−6^, CD8^+^: *p*=6.3×10^−10^ by Mann-Whitney U test) than those associations found with the single chains individually (Fig. 2H-I). Similarly, the association between J*αβ* pairs was significantly stronger (CD4^+^: *p*=9.8×10^−7^, CD8^+^: *p*=2.1×10^−4^ by Mann-Whitney U test) than those of either the *α* or *β* chain alone (Sup. Fig. 8H-I).

Biologically, this finding is consistent with the notion that both the *α* and *β* chain contribute substantially to TCR-pMHC binding^26–32^. These findings additionally highlights the importance of new single-cell methods that allow for the capture of paired *αβ* chains over traditional bulk-sequencing methods that allow only for the capture of individual chains.

### CDR3 features are weakly associated with T-cell lineage

The TCR-pMHC interaction is also dependent upon the contributions of the CDR3 regions of both the *α* and *β* chains^26–32^, leading us to investigate the relationship between CDR3 sequence and T-cell lineage. Examining the frequency with which each amino acid occurred across the single-chain CDR3 repertoires shows strong differences between the *α* and *β* chains (Fig. 3A-D). This is likely due to the differences in amino acid usage in the *α* and *β* chain V(D)J germline regions. However, we observed only small differences in amino acid use between the CD4^+^ and CD8^+^ repertoires (Fig. 3E-F). Previous studies have observed an association between CDR3 net charge and T-cell lineage^38,39^, consistent with our findings that net CDR3 charge, but not CDR3 length, is associated with T-cell lineage for both the *α* and *β* chains (Fig. 3G-I and Sup. Fig. 12A-C).

**Figure 3:**
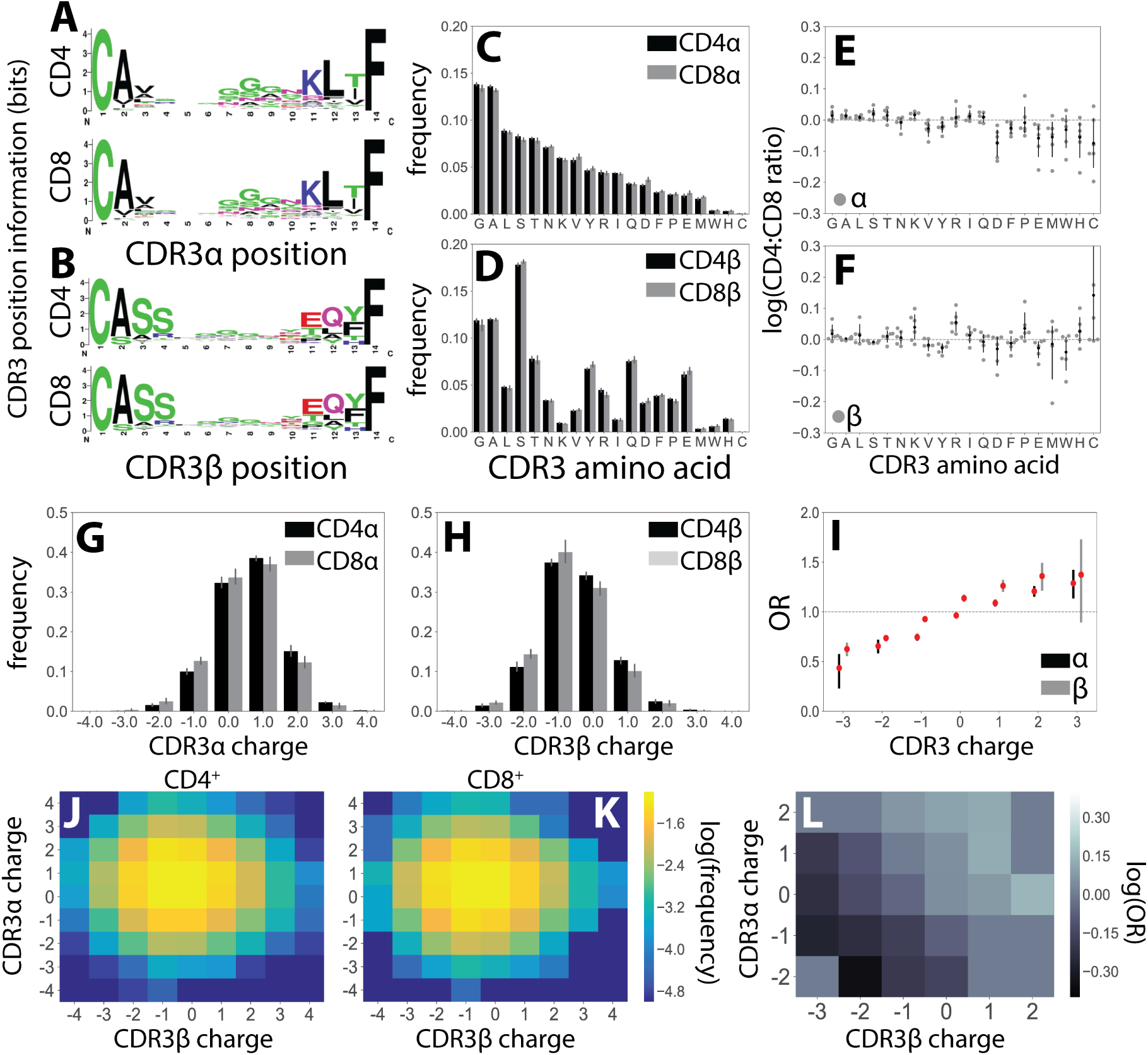
CDR3 features correlate weakly with T-cell lineage. WebLogos ^44^ show composite sequence information for **(A)** *α* and **(B)** *β* chains of length 14 amino acids for the CD4^+^ (top) and CD8^+^ (bottom) T-cell lineages. **(C)** Usage frequencies for all 20 amino acids, rank ordered by prevalence in CDR3*α*, are shown for CDR3*α* and **(D)** CDR3*β* sequences across the CD4^+^ and CD8^+^ repertoires. **(E)** The CD4^+^:CD8^+^ usage ratio for all amino acids are shown for the *α* and **(F)** *β* chains. The frequency with which each amino acid is used is shown for each individual (gray circles) with the population mean and standard deviation shown in black. **(G)** Single chain net charge distributions for both CD4^+^ and CD8^+^ TCR repertoires are shown for the predominately positively charged *α* chains and **(H)** the predominately negatively charged *β* chains. **(I)** Odds ratios (OR) quantify the strength of association of CDR3 net charge with lineage for both the *α* (black) and *β* (gray) chains. Red markers indicate statistical significance (*p<*0.05 after Bonferonni correction). **(J)** The log frequency for each CDR3*αβ* charge pair is shown for the CD4^+^ and **(K)** CD8^+^ TCR repertoires. **(L)** Significant (*p<*0.05 after Bonferroni correction) log odds ratios reveals strong CD4^+^:CD8^+^ bias for 21 CDR3*αβ* charge pairs.

We further examined the relationship of paired CDR3*αβ* charge and length with T-cell lineage (Fig. 3J-K and Sup. Fig. 12D-E). Again calculating the odds ratio, we found 21 CDR3*αβ* charge pairs and 14 CDR3*αβ* length pairs associated with a significant CD4^+^:CD8^+^ bias (Fig. 3L and Sup. Fig. 12F). We additionally observe that paired *αβ* chain lengths tend to be associated with stronger biases towards CD4^+^ status than either of the single chains alone (Sup. Figs. 12G). Surprisingly, however, no significant differences were observed in the strength of association between paired and single-chain CDR3 length for CD8^+^ status or for CDR3 charge for either CD4^+^ or CD8^+^ status (Sup. Figs. 12H and 13).

### Paired chain sequences are more informative of CD4^+^-CD8^+^ status than single chains

In order to better understand the amount of information about CD4^+^ and CD8^+^ status encoded in the *α*, *β*, and *αβ* TCR sequences, we next quantified the mutual information^33,45^, corrected for finite sample sizes, between several TCR features and T-cell lineage (Table 1). Examining V and J usage, as well as CDR3 length, we find that paired sequences carry more information about lineage than either of the single chains alone. Particularly for V*αβ*, we observe synergistic information^46^ in which the paired chains carry more information than the individual chains summed together.

**Table 1:** Mutual information between TCR features and T-cell lineage. Mutual information estimates (bits) were calculated using a finite-sampling correction to quantify the amount of information about T-cell lineage by various TCR features drawn from the *α*, *β*, and paired *αβ* repertoires.

| | $\alpha$ | $\beta$ | $\alpha\beta$ |
| --- | --- | --- | --- |
| **V** | 0.015 | 0.013 | 0.035 |
| **J** | 0.005 | 0.001 | 0.006 |
| **CDR3 Charge** | 0.003 | 0.004 | 0.003 |
| **CDR3 Length** | 0.002 | 0.001 | 0.005 |

We next investigated whether the use of paired sequences would better allow us to predict T-cell lineage from TCR features using machine learning classifiers. Using a multi-layer perceptron neural network classifier, we demonstrate that the *α* and *β* chain are both weakly informative of lineage and that paired TCR sequences carry substantially more information than either the *α* chain (*p*=9.0×10^−5^) or *β* chain (*p*=9.1×10^−5^ by Mann-Whitney U test) alone (Fig 4). Similar results were obtained using both support vector machine (SVM) and logistic regression classifiers (Sup. Fig. 14) ^38,39^. From a biological perspective, this finding is consistent with a mechanistic model in which both chains contribute to the TCR-pMHC interaction.

**Figure 4:**
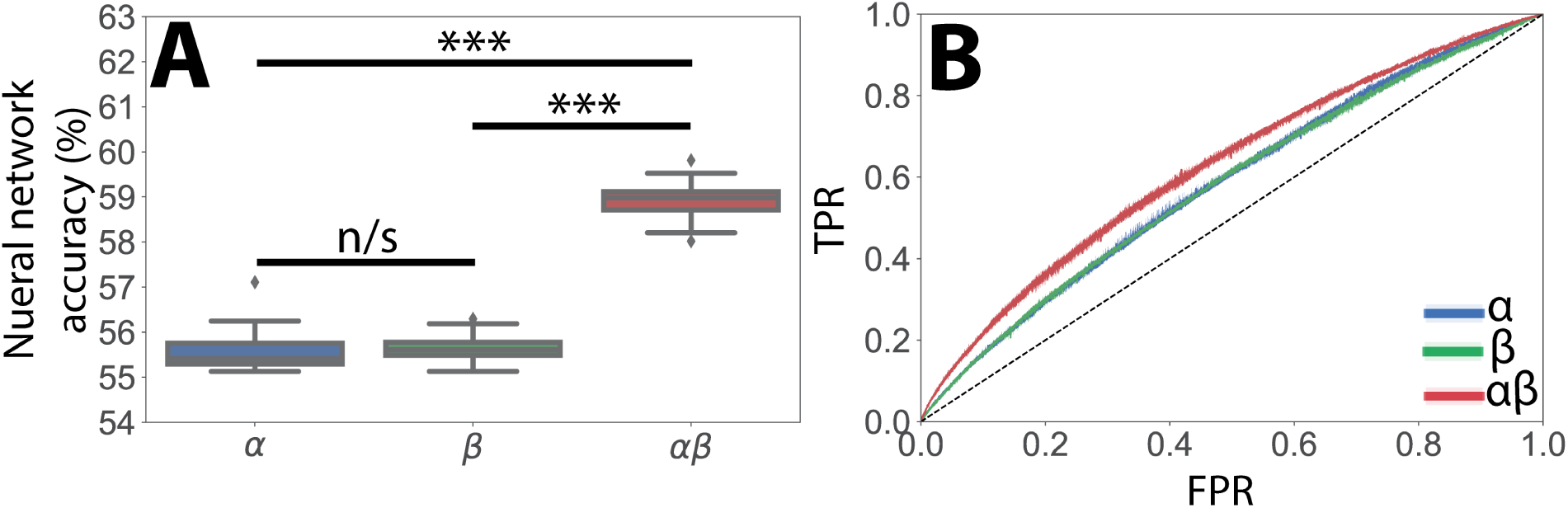
Paired *αβ* sequences are more informative of T-cell lineage than single chain sequences alone. **(A)** A multi-layer perceptron (MLP) neural network classifier was trained using a constant length vector encoding V and J region usage, CDR3 length, and CDR3 amino acid usage frequencies for the *α*, *β* and *αβ* repertoires. Classifiers trained using paired *αβ* TCR sequences substantially outperformed those trained using either the *α* or *β* chain alone. Boxplots show accuracy in predicting CD4^+^ or CD8^+^ from *α*, *β* or *αβ* TCR information through 10 rounds of bootstrapping. Statistical significance between groups was assessed by Mann-Whitney U test (\*\*\**p<*0.001, n/s-not significant at *p <* 0.05 level). **(B)** Receiver operator characteristic (ROC) curves showing true positive rate (TPR) versus false positive rate (FPR) for a neural network classifier trained on *α*, *β* and *αβ* TCR sequences again show models using paired *αβ* sequence information outperform those trained on only a single chain.

Of note is a previous report using a SVM classifier and CDR3 length-dependent parametrization to predict T-cell lineage from TCR sequences with greater than 90% accuracy ^39^. This approach, however, failed to achieve the same degree of predictive accuracy when using our dataset (Sup. Fig. 15). To better understand this finding, we compared the TCR sequences from this study ^39^ with those reported here and an additional bulk-sequencing TCR*β* dataset^40^. We find that the aforementioned increased predictive accuracy is driven by anomalous V*β* and J*β* gene frequencies in the Li *et al.* dataset, possibly due to a lack of rigorous PCR correction, as compared with the other two datasets (Sup. Figs 16-18).

## 3. Conclusions

In summary, we have created the largest database of paired *αβ* TCR sequences to date. Our analysis of the healthy CD4^+^ and CD8^+^ TCR repertoires revealed systematic differences between the two T-cell populations, particularly in the utilization of V*αβ* pairings. Furthermore, we have presented one of the first comprehensive analyses of the *α* chain repertoire, showing both chains are similarly informative of T-cell lineage. Finally, utilizing approaches from information theory and machine learning, we have shown that features of the paired *αβ* TCR are substantially more informative of lineage than individual chains. Our results thus provide new evidence for the role of germline-encoded TCR-pMHC interactions and implicate both chains as playing important roles in determining TCR interactions. We believe that the rigorous examination of the normal TCR repertoires presented in this study both demonstrates the utility of capturing *αβ* pairs in profiling the TCR repertoire and will prove to be valuable in understanding the perturbations caused by infectious, oncological and auto-immune disease states^47–53^.

## 4. Materials and Methods

### Single-cell barcoding and sequencing

TCR sequences for subjects 1-5 were obtained from Grigaityte *et al.*^33^ In brief, peripheral blood mononuclear cells (PBMCs) were obtained from five healthy donors after appropriate informed consent. Blood samples then underwent a pan T-cell enrichment, were tagged with unique barcodes *via* a newly developed single-cell barcoding in emulsion technology^24^, and sequenced using an Illumina MiSeq sequencer. Raw sequences were processed using a custom pipeline^33^ to identify *αβ* pairs utilizing MiXCR 2.2.1^54^ to identify V(D)J segments and annotate the CDR3 region of each TCR.

TCR sequences for Subject 6 were similarly obtained from a commercially purchased PBMC sample (ATCC PCS-800-011TM) drawn from a healthy individual. CD4^+^ and CD8^+^ T-cell populations were separated using magnetic bead enrichment according to the manufacturer protocol (EasySep Human T Cell Enrichment Kit, StemCell Technologies). The PBMC samples used in Grigaityte *et al.*^33^ for S1 and S3 were additionally obtained and sorted into CD4^+^ and CD8^+^ using fluorescence activated cell sorting (Becton Dickinson FACSARIA SORP). For these samples, cells were barcoded in emulsion^25^ using the Chromium Controller using the Single Cell V(D)J reagent kit (10X Genomics) and sequenced using an Illumina HiSeq 2500 sequencer. Raw sequencing reads were processed using the computational pipeline previously described^33^.

The Li *et al.* dataset^39^ was provided by N.P. Weng as a processed datafile containing VJ segments and CDR3 amino acid sequences. The Emerson *et al.* dataset^40^ was downloaded from Adaptive Biotechnologies open-access immuneACCESS database (https://clients.adaptivebiotech.com/immuneaccess). Of note, though the original study consisted of both TCR sequences obtained from healthy and disease patients, only the 17 healthy samples are used here.

#### Data analysis

Following the processing described above, we generated text files containing information about V(D)J segment use and CDR3 nucleotide and amino acid sequence for each of the identified paired *αβ* TCR sequences (Supplemental Figures 1 and 2). As we care about identifying features of the TCR repertoires between the CD4^+^ and CD8^+^ populations, we count each unique TCR clonotype only once. That is, clonal expansion of random clones in the CD4^+^ and CD8^+^ would bias our analysis of the factors that effect differentiation. As such, we include each TCR clonotype only once into our final dataset. Here, we define a clonotype to be the V*αβ* regions used and amino acid CDR3*αβ* sequences. We then identified TCR clonotypes that were shared between the CD4^+^ and CD8^+^ compartments.

The degree of overlap between the CD4^+^ and CD8^+^ TCR repertoires was quantified using the Jaccard Index (*J*):

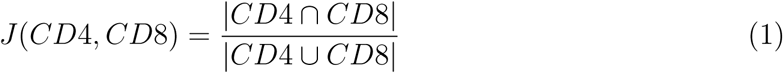

Here |*CD*4∩*CD*8| refers to the cardinality of the intersection between the CD4^+^ and CD8^+^ TCR repertoires (*i.e.* the number of TCRs found in both repertoires). |*CD*4 ∪ *CD*8| refers to the union of the two repertoires (*i.e.* the number of TCRs found in either of the two repertoires). The Jaccard Index was calculated independently for the *α* (*J*(*CD*4_*α*_, *CD*8_*α*_)), *β*(*J*(*CD*4_*β*_, *CD*_8_*β*)), and *αβ* (*J*(*CD*4_*αβ*_, *CD*_8_αβ)) TCR repertoires. TCR sequences shared between the CD4^+^ and CD8^+^ TCR repertoires were excluded from the machine learning classification analysis.

Furthermore, as done previously^33^, the paired *αβ* repertoire consists of all unique, paired TCR sequences and the *α* and *β* individual chain repertoires were derived directly from the paired repertoire. That is, the individual *α* repertoire consists of all the *α* chains present in the paired dataset. Thus, the *α*, *β*, and *αβ* datasets are all of the same size and differences in sample size do not drive the observed differences. Furthermore, all boxplots represent median and inter-quartile range.

All analysis steps, unless otherwise noted, were performed using custom Python scripts available at our Github repository (https://github.com/JasonACarter/CD4CD8-Mansucript).

#### VJ segment usage

V(D)J segments were identified from raw sequences by MiXCR and annotated according to the International ImMunoGeneTics (IMGT) V(D)J gene definitions^55^. The odds ratio (OR) for a given TCR characteristic and T-cell lineage was calculated by counting the number of TCRs with (*C*^+^) and without (*C^−^*) that characteristic within the CD4^+^ (*T* ^4^) and CD8^+^ (*T* ^8^) repertoires. The OR is then given as:

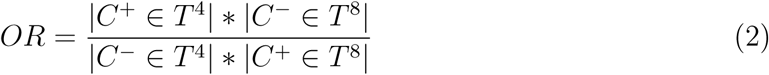

That is, the numerator is the number of CD4^+^ TCRs with a given feature are multiplied by the number of CD8^+^ TCRs without that feature. The denominator is given by the number of CD4^+^ cells without that feature multiplied by the number of CD8^+^ with that feature. Thus, an OR greater than 1 corresponds with a bias towards CD4^+^ and an OR less than 1 corresponds with a CD8^+^ bias. 95% confidence intervals and a p-value were then calculated for each OR using Fisher’s exact test implemented using the SciPy library (www.scipy.org). Multiple hypothesis testing correction was applied to single chain *p*-values using a Bonferroni correction and paired chains p-values were converted to *q*-values^56^. Significance was assessed at the *p<*0.05 or *q<*0.05 level.

#### CDR3 features

Sequence logos showing the amino acid frequency for a given position in the sequence were generated using all *α* and *β* CDR3 sequences of length 14 using WebLogo^44^. Of note, we defined the CDR3 length to be inclusive of the proximal cysteine and terminal phenylalanine that define the CDR3 region. The ratio of each amino acid in CDR3 between the CD4^+^ and CD8^+^ populations was calculated by dividing the frequency of a given amino acid across all CD4^+^ CDR3 sequences for a given chain by the frequency with which that amino acid occurred across all CD8^+^ CDR3 sequences. CDR3 charge was calculated as the sum of negatively charged amino acids (D and E) and positively charged amino acids (R and K) present in the CDR3 region.

#### Mutual information

The mutual information^45^ (I), in bits, between a given feature, X, and T-cell lineage (L) was calculated as:

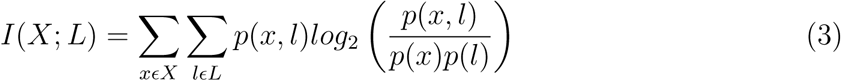

In order to correct for biases in our MI estimate arising from our limited sample sizes, we then applied a bootstrapping based finite-sampling correction previously described^33,57^. We additionally calculate the synergistic information ^46^ (*S*) according to:

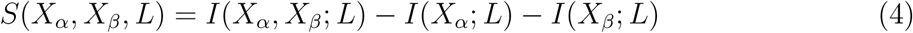

where *X_α_* and *X_β_* refer to TCR*α* and TCR*β* features, respectively.

#### Machine learning

Multi-layer perceptron (MLP) neural network, logistic regression, and support vector machine (SVM) classifiers were implemented using custom Python scripts employing sklearn’s SVM library^58^. For SVM’s trained on the Li *et al.* and Emerson *et al.* dataset, CDR3*β* amino acid sequences were first converted in numeric vectors using Atchley factors^39,59^. As the length of these numeric vectors depended on the length of the CDR3 region, a separate SVM was trained for each CDR3 length between 10 and 15. For all machine learning classifiers, each dataset was divided into a training set (75%) and a testing set (25%) and the accuracy of the testing set was reported for both the CD4^+^ and CD8^+^ populations. Standard deviations were calculated *via* 10 rounds of bootstrapping.

For our dataset, we wished to understand if the paired *αβ* repertoire was more informative than either of the single chain repertoires. As converting each CDR3*αβ* pair into a numeric vector would drastically lower our sample size, we developed a new methodology for preparing input vectors for TCRs that are independent of the CDR3 length. Specifically, we designated a TCR’s V and J segment as categorical variables. Additionally, we included the length of each CDR3 region and the frequency of each of the twenty amino acids used in the CDR3 region. Although this methodology loses information encoded in the amino acid sequence of the CDR3 region, it still captures many of the salient features we find to carry information about T-cell lineage and has the advantage of not quickly diminishing our sample size as a length-dependent method would.

## 5. Acknowledgments

The authors thank Doug Fearon for comments on the manuscript, Pamela Moody and the CSHL Flow Cytometry Shared Resource for help with FACS experiments and the CSHL DNA Sequencing Core for next-generation sequencing. We additionally thank N.P. Weng for providing the *β* chain bulk sequencing dataset from Li *et al.* JAC was partially supported by NIHGM MSTP Training award T32-GM008444 and a LIBH grant. KG was funded by the Ferish-Gerry fellowship from the Watson School of Biological Sciences. GA was funded by the Simons Foundation and the Stand Up To Cancer-Breast Cancer Research Foundation Convergence Team Translational Cancer Research Grant, Grant Number SU2C-BCRF 2015-001.

---

**Supplemental Figure 1:**
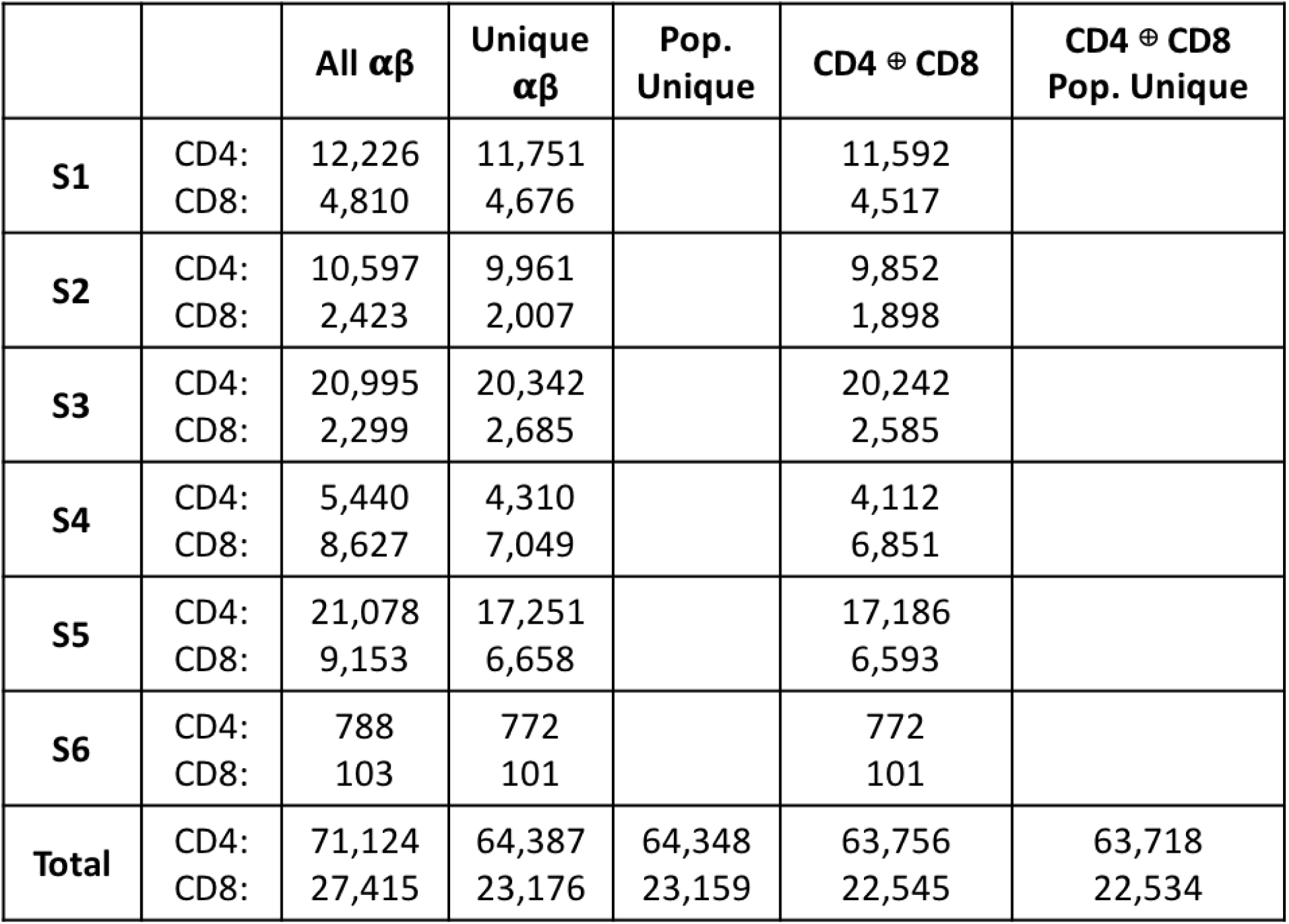
Table showing number of paired sequences at each processing step for each individual. Peripheral blood mononuclear cells (PBMCs) were previously obtained from 5 healthy individuals (S1-S5) and sequenced using single-cell barcoding in emulsion^24,33^. The original PBMC samples from S1 and S3 (see Sup. Fig.2), as well as a new sample from an additional healthy individual (S6), were sequenced using a commercially available single-cell system (10x Genomics) ^25^. In all, we obtain 71,124CD4^+^ and 27,415 CD8^+^ cells with productive V(D)J rearrangements in both chains (all *αβ*). For each individual, we then assessed the set of unique *αβ* clonotypes defined by V*αβ* and CDR3*αβ* sequences (unique CDR3*αβ*). From the set of unique clonotypes we then removed any sequences found both in the CD4^+^ and CD8^+^ repertoires (individual CD4^+^ ⊕ [exclusive or] CD8^+^). The set of unique clonotypes found only in the CD4^+^ or CD8^+^ repertoires between all individuals was then assembled (population unique).

**Supplemental Figure 2:**
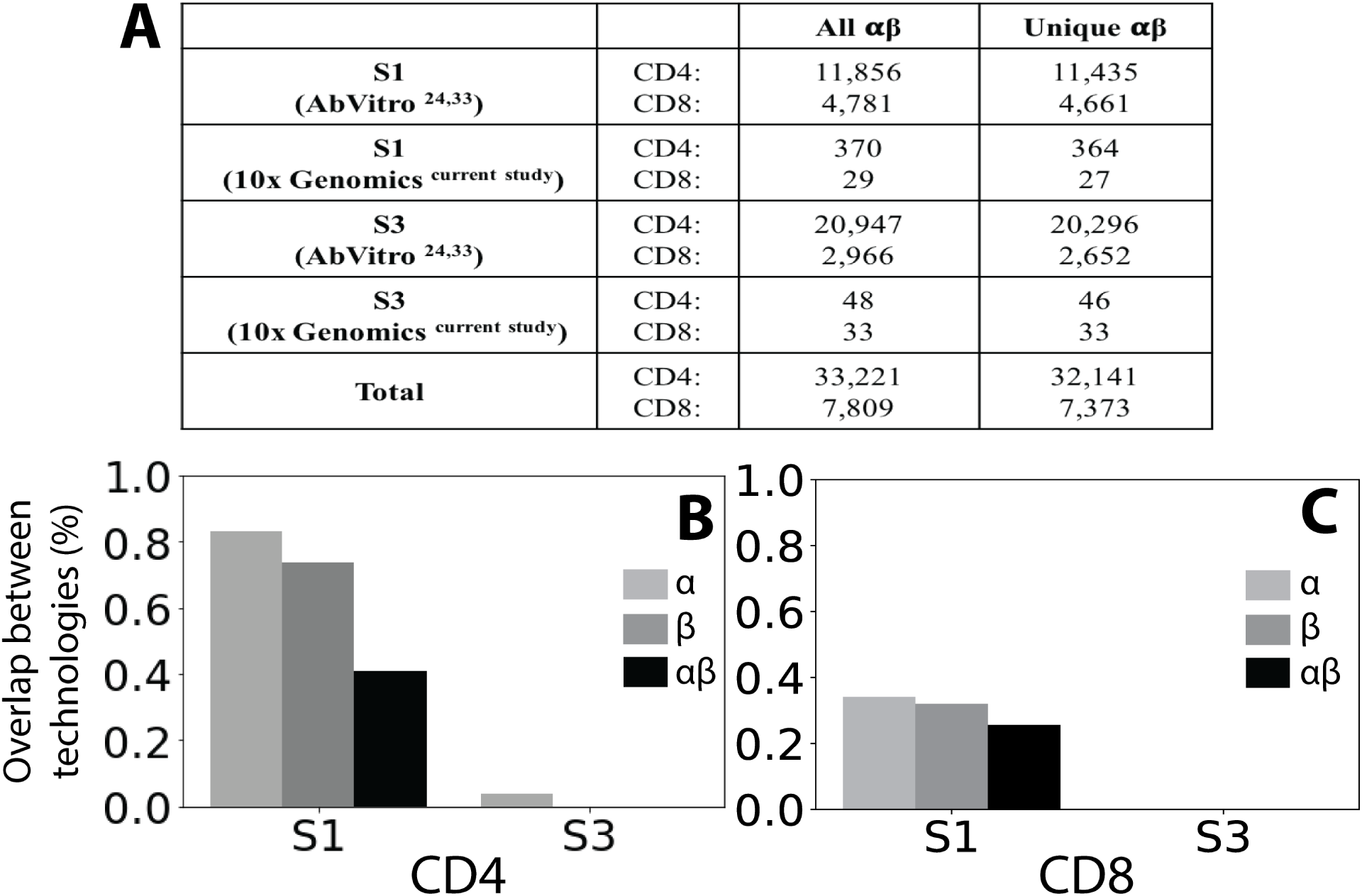
Comparison of individual samples sequenced independently with two single-cell technologies. Samples from subjects 1 and 3 (S1 and S3) were sequenced previously ^33^ using a novel single-cell emulsion barcoding strategy (AbVitro)^24^. We additionally resequenced samples from these subjects using a commercially available single-cell sequencing set-up (10x Genomics) ^25^. **(A)** We report the total number of productive *αβ* TCR pairs (All *αβ*) and the number of unique TCRs per sample (Unique *αβ*). **(B)** For each subject, we then assessed the number of *α*, *β*, and paired *αβ* TCR sequences observed in both sequencing replicates for the CD4^+^ and **(C)** CD8^+^ populations.

**Supplemental Figure 3:**
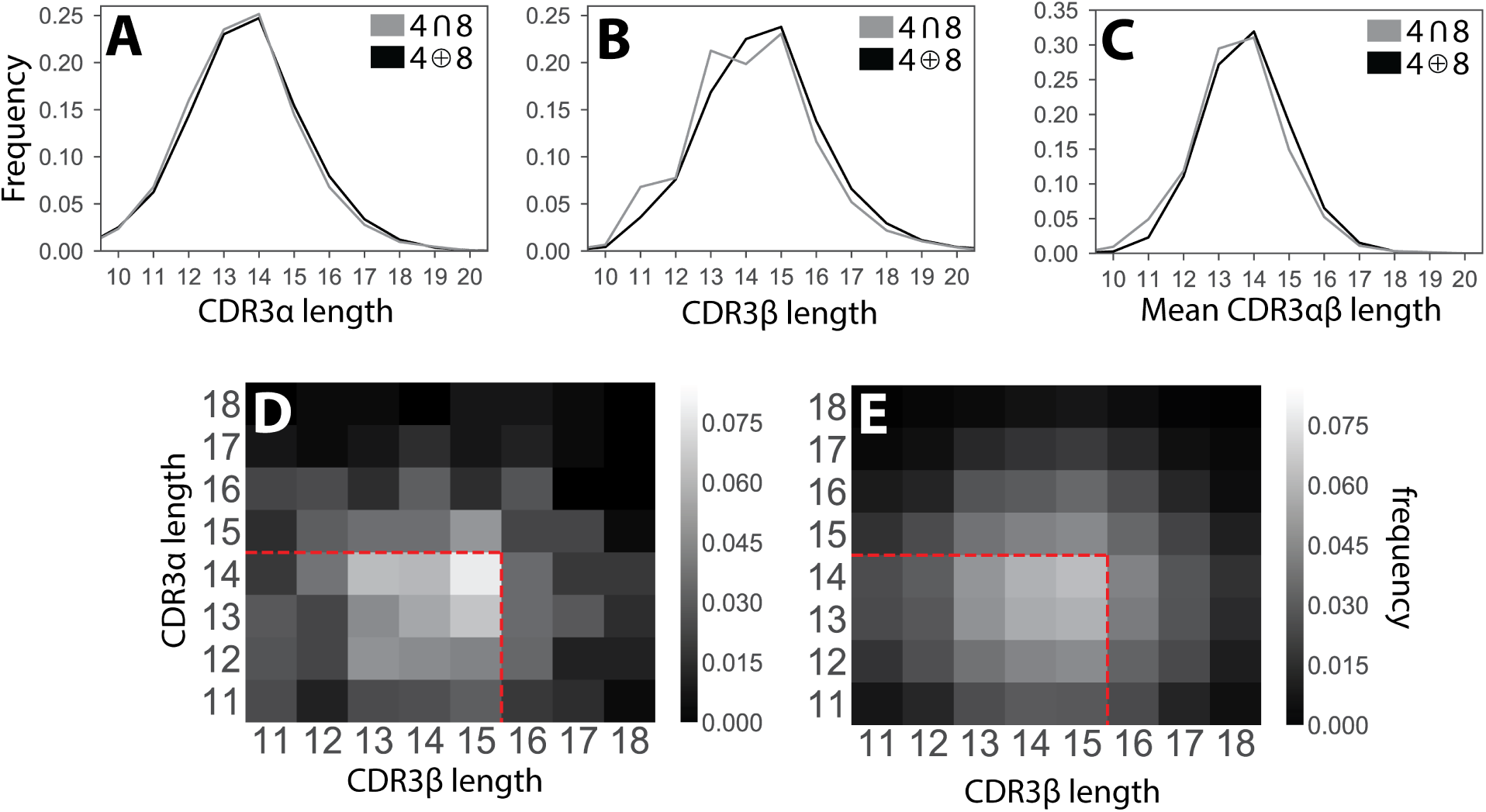
CDR3 sequences shared between the CD4^+^ and CD8^+^ repertoires tend to be shorter than those found in only one repertoire. CDR3 length distributions show sequences found in both the CD4^+^ and CD8^+^ repertoires (∩) are shorter than those found in only one of the two repertoires (⊕) for the **(A)** *α*, **(B)** *β*, and **(C)** paired *αβ* repertoires. For paired sequences, we report the average length of the *α* and *β* chains.**(D)** Heatmaps showing frequency with which each *α* and *β* CDR3 length pair is present in the TCR repertoire shared between the CD4^+^ and CD8^+^ lineages and for the **(E)** TCR repertoire present in only one of the two lineages. Dashed red lines indicate the average length for the *α* (14 amino acids) and *β* chains (15 amino acids).

**Supplemental Figure 4:**
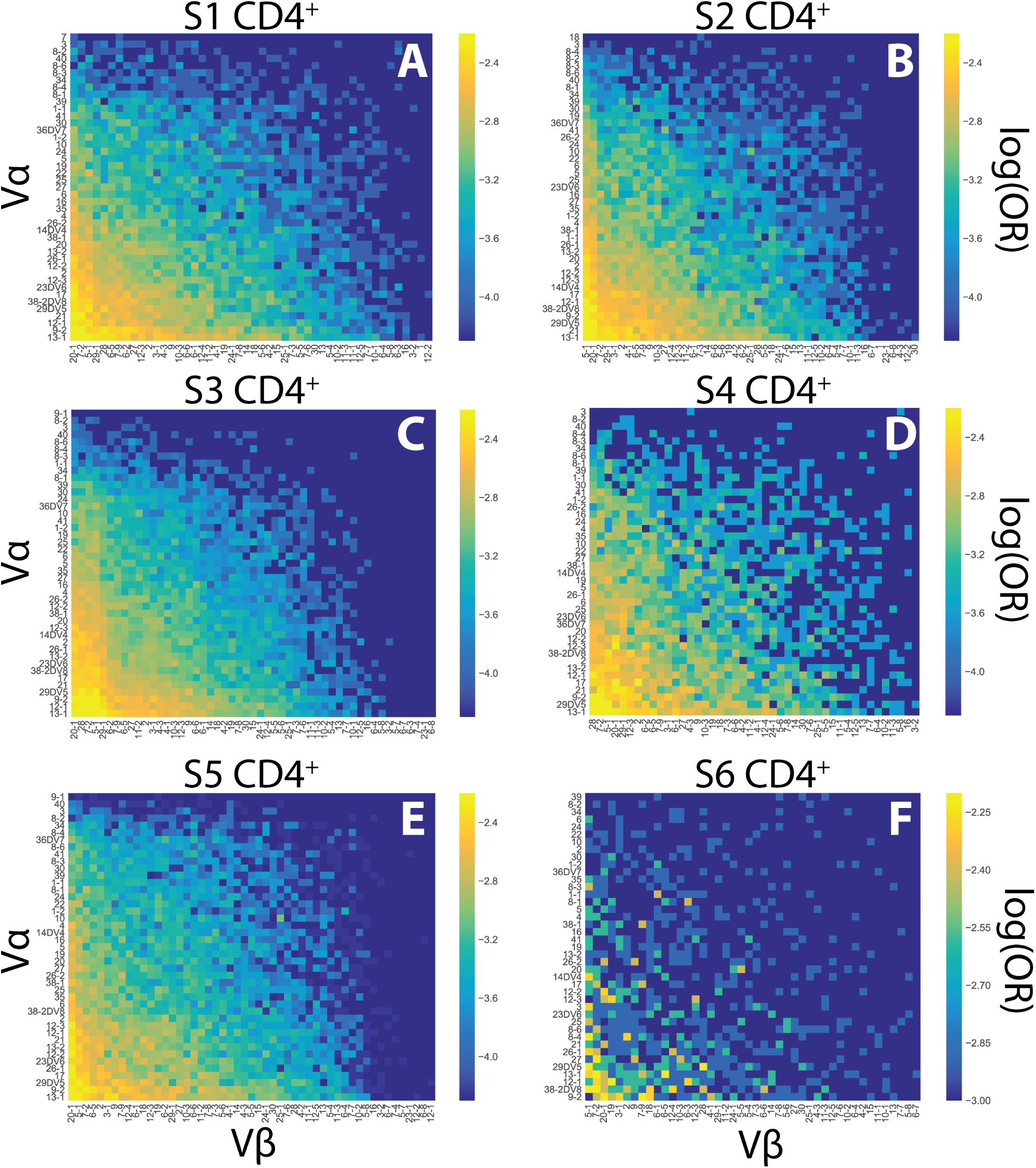
V*αβ* usage in the CD4^+^ TCR repertoire for each individual. Log frequency heatmaps show distribution of V*αβ* pairs within the CD4^+^ TCR repertoire for each individual. **(A)** S1 (n=11,751), **(B)** S2 (n=9,961), **(C)** S3 (n=20,242), **(D)** S4 (n=4,310), **(E)** S5 (n=17,251), **(F)** S6 (n=722)

**Supplemental Figure 5:**
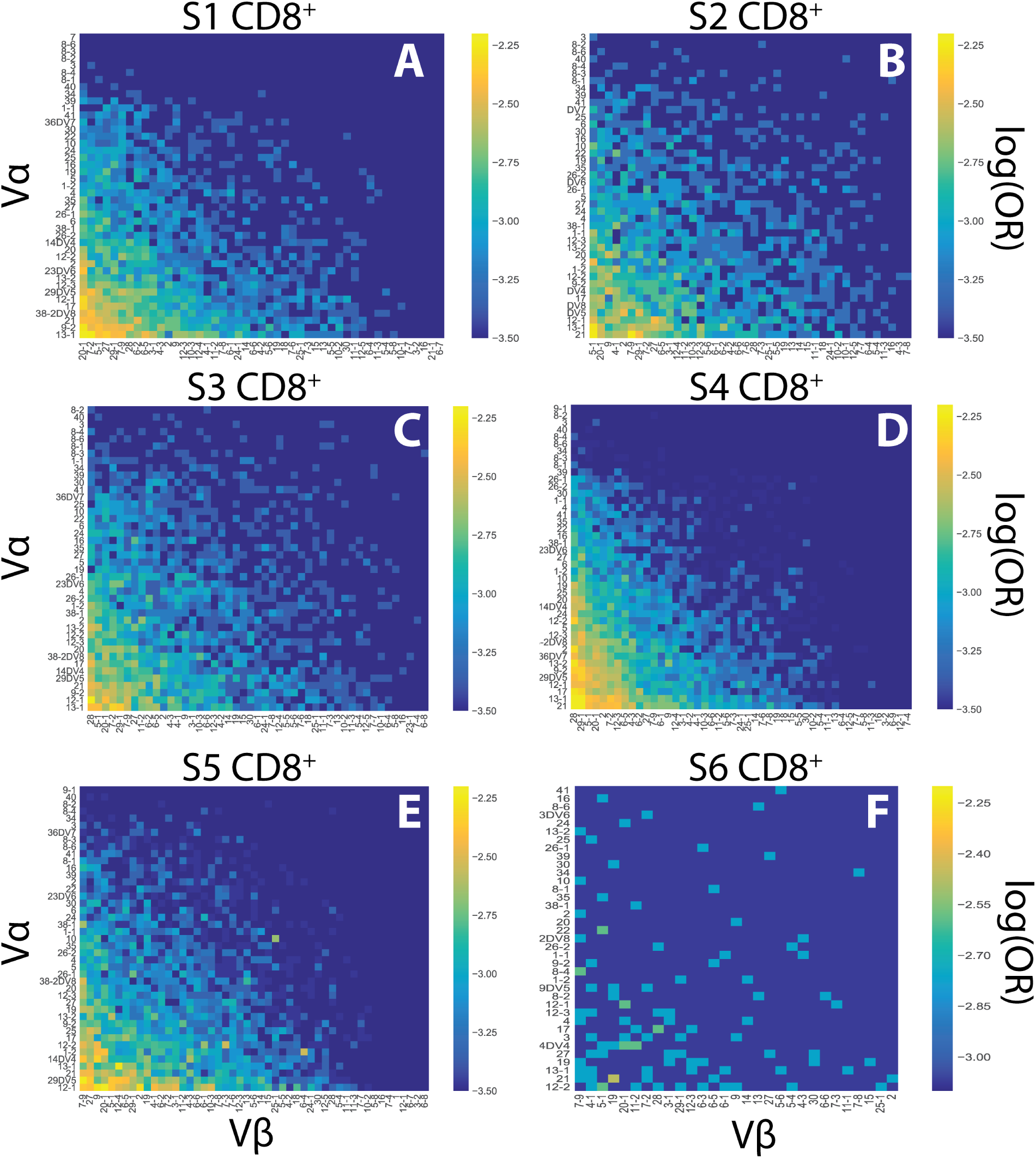
**V***αβ* usage in the CD8^+^ TCR repertoire for each individual. Log frequency heatmaps show distribution of V*αβ* pairs within the CD8^+^ TCR repertoire for each individual. **(A)** S1 (n=4,676), **(B)** S2 (n=2,007), **(C)** S3 (n=2,685), **(D)** S4 (n=7,049), **(E)** S5 (n=6,658), **(F)** S6 (n=101)

**Supplemental Figure 6:**
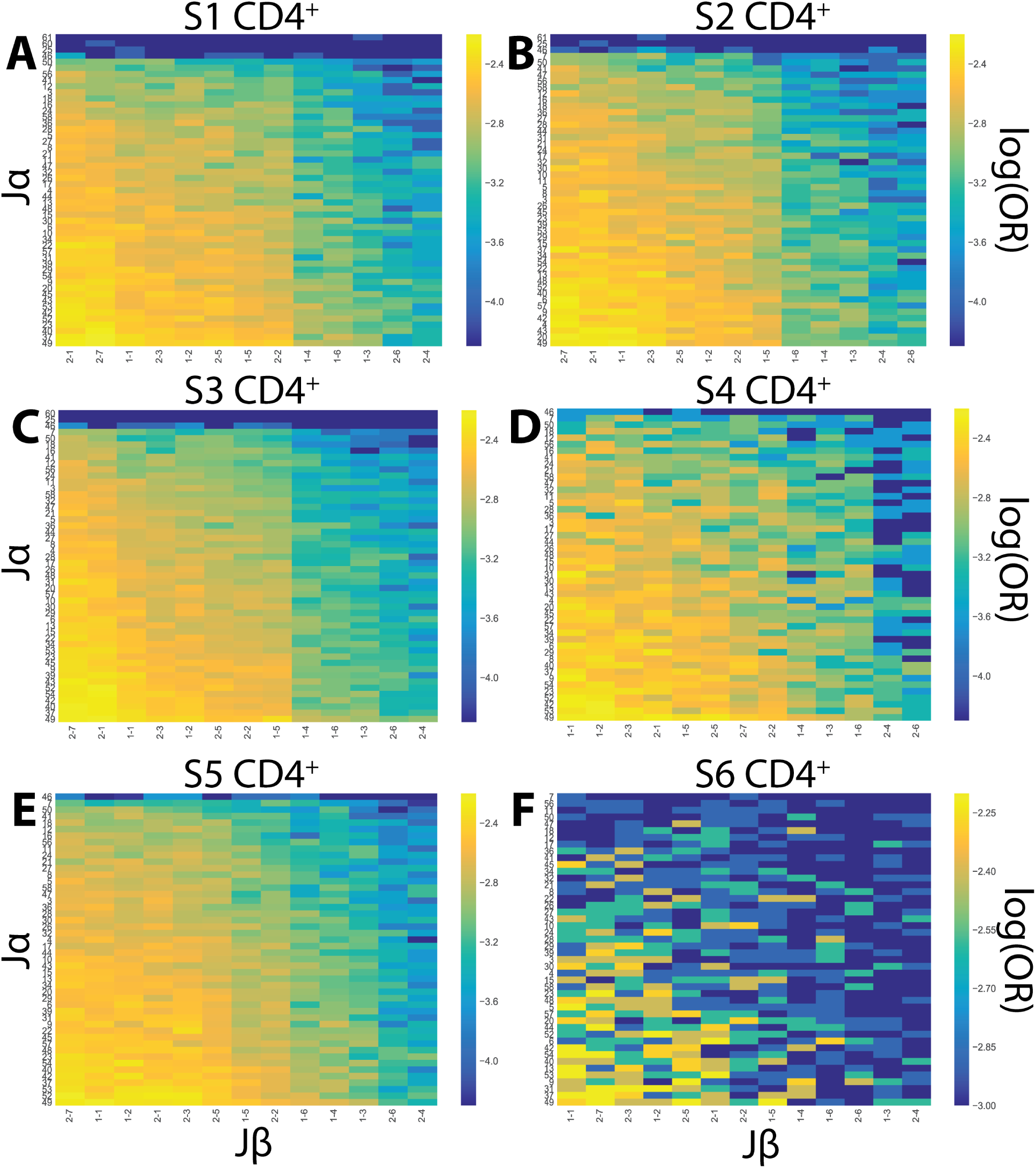
J*αβ* usage in the CD4^+^ TCR repertoire for each individual. Log frequency heatmaps show distribution of J*αβ* pairs within the CD4^+^ TCR repertoire for each individual.**(A)** S1 (n=11,751), **(B)** S2 (n=9,961), **(C)** S3 (n=20,242), **(D)** S4 (n=4,310), **(E)** S5 (n=17,251), **(F)** S6 (n=722)

**Supplemental Figure 7:**
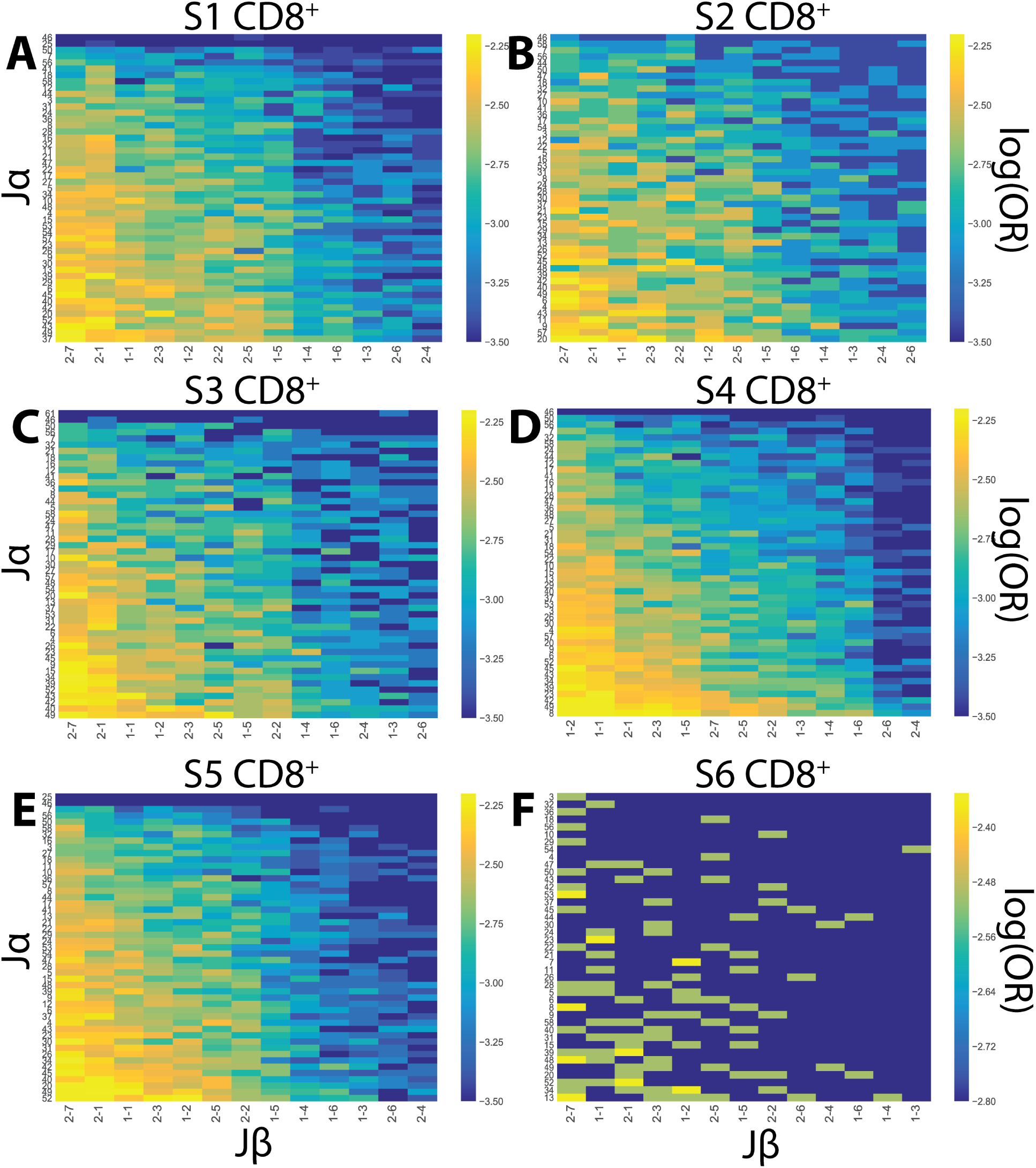
J*αβ* usage in the CD8^+^ TCR repertoire for each individual. Log frequency heatmaps show distribution of J*αβ* pairs within the CD8^+^ TCR repertoire for each individual. **(A)** S1 (n=4,676), **(B)** S2 (n=2,007), **(C)** S3 (n=2,685), **(D)** S4 (n=7,049), **(E)** S5 (n=6,658), **(F)** S6 (n=101)

**Supplemental Figure 8:**
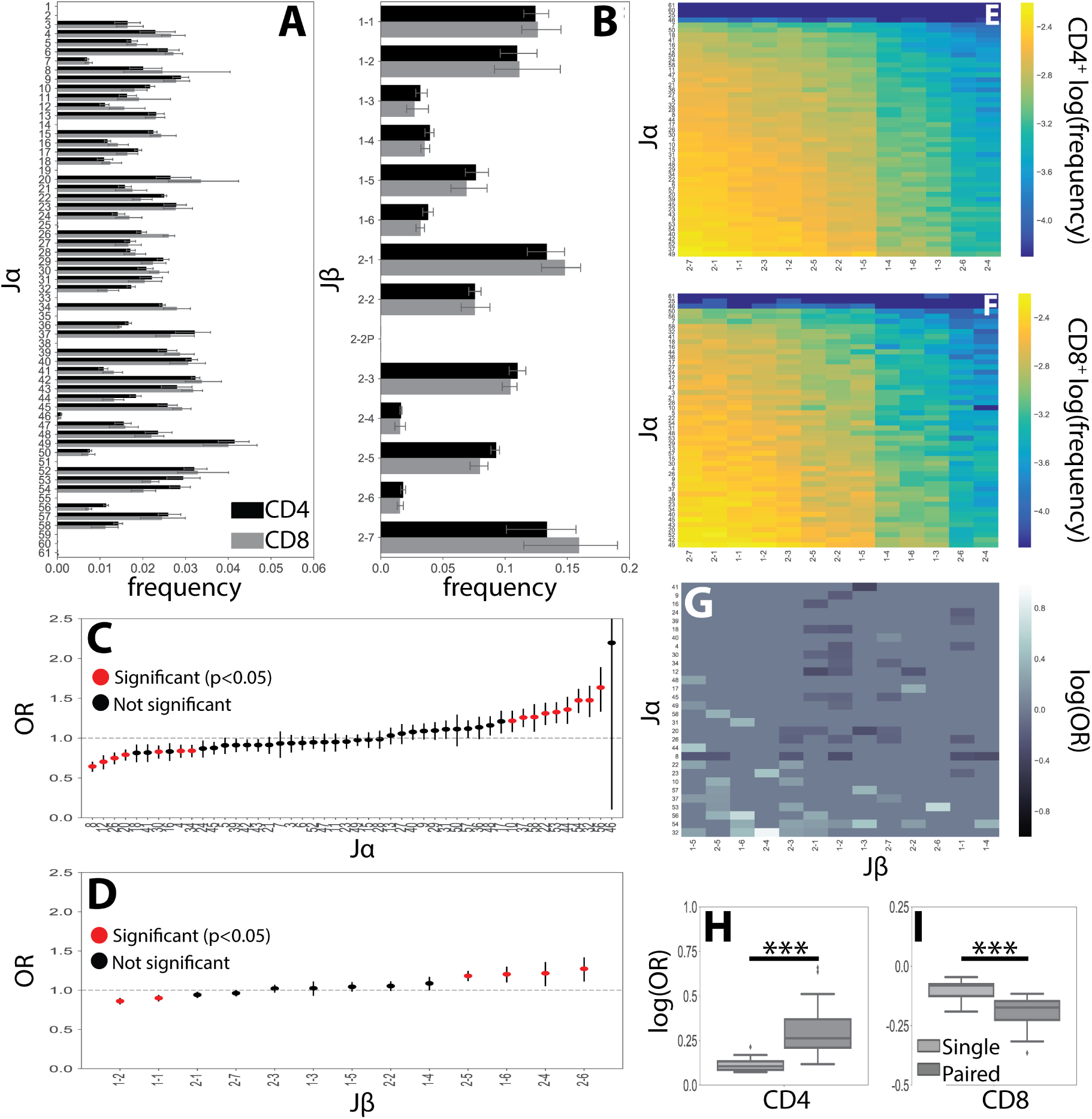
J germline region usage in the *α*, *β* **and** *αβ* repertoires. **(A)** J*α* and **(B)** J*β* single-chain germline region usage frequencies were calculated for each individual’s CD4^+^ and CD8^+^ T-cell lineages. Error bars represent the standard deviation across individuals. **(C)** The CD4^+^ (n=63,718) and CD8^+^ (n=22,534) TCR repertoires were then pooled across individuals and the CD4^+^:CD8^+^ odds ratio (OR) was calculated for each J*α* and **(D)**J*β* germline region. An OR*>* 1 represents a CD4^+^ bias, while an OR*<* 1 represents a CD8^+^ bias with error bars representing the 95% confidence interval. The mean is represented by a red or black dot, with red representing statistical significance at the *p<*0.05 by Fisher’s exact test level after applying Bonferroni correction. **(E)** Paired J*αβ* usage frequencies across all individuals for CD4^+^ and **(F)** CD8^+^ TCR repertoires. **(G)** Significant (*q<*0.05 by Fisher’s exact test) log odds ratios reveals strong CD4^+^:CD8^+^ biases for 72 J*αβ* pairs. **(H)** Boxplots were calculated for the set of all significant odds ratios associated with single chains (J*α* or J*β*) and compared with those associated with J*αβ* pairs. Paired associations for both CD4^+^ and **(I)** CD8^+^ status were significantly stronger (\*\*\**p<* 0.001 by Mann-Whitney U test) than those associated with a single chain alone.

**Supplemental Figure 9:**
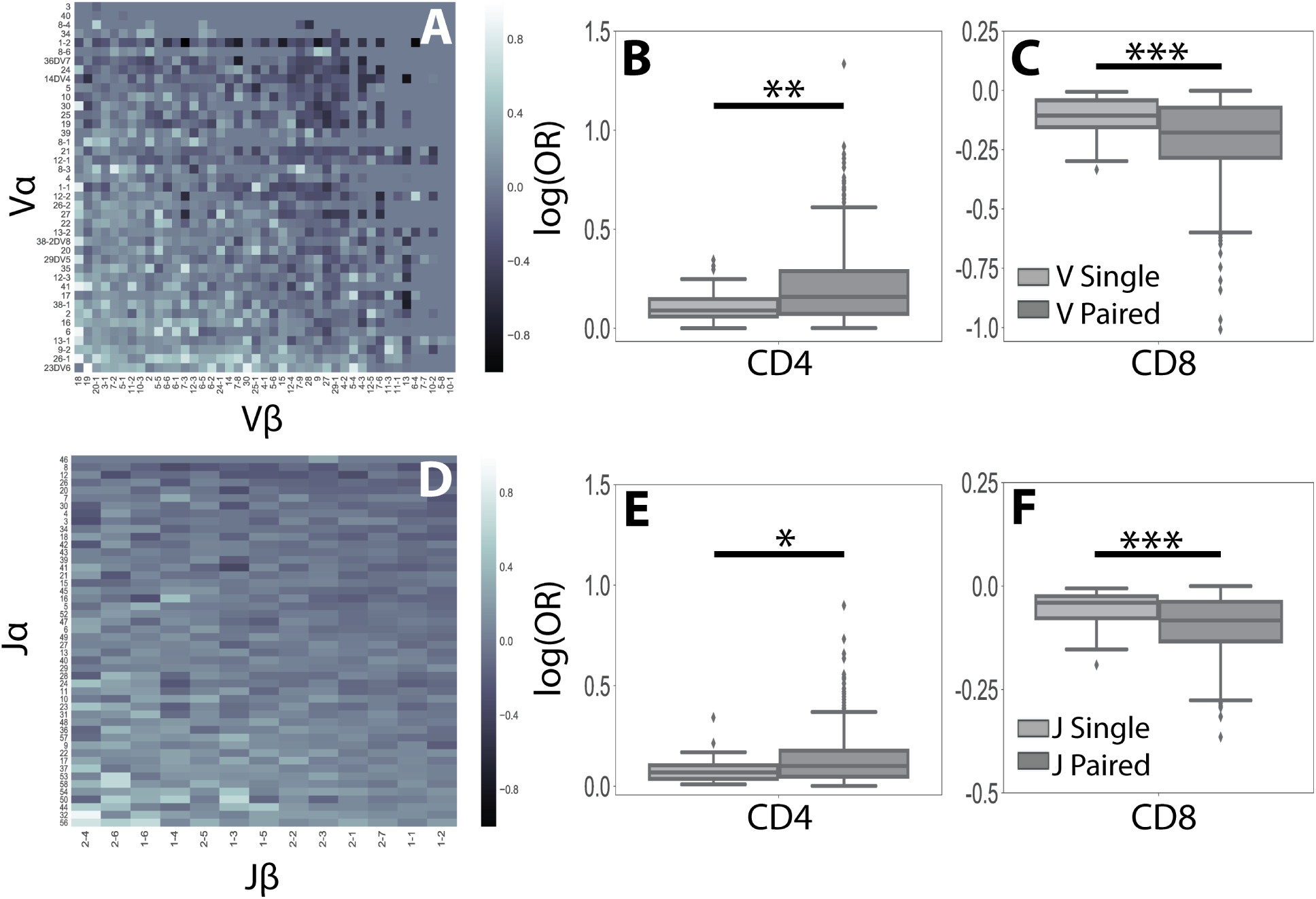
Odds ratios calculated for all V*αβ* **and J***αβ* pairs. Log odds ratios were calculated for all **(A)** V*αβ* pairs and **(D)** J*αβ* pairs. Boxplots showing that all odds ratios for V*αβ* pairs are more strongly associated with T-cell lineage than either single chain alone for **(B)** CD4^+^ status (*p*=1.5×10^−3^) and **(C)** CD8^+^ status (*p*=2.6×10^−5^). **(E)** Similarly, J*αβ* pairs are associated with stronger CD4^+^ (*p*=1.1×10^−2^) and **(F)** CD8^+^ biases (*p*=5.7×10^−4^) than either of the single chains alone. All *p* values are obtained by Mann-Whitney U test.

**Supplemental Figure 10:**
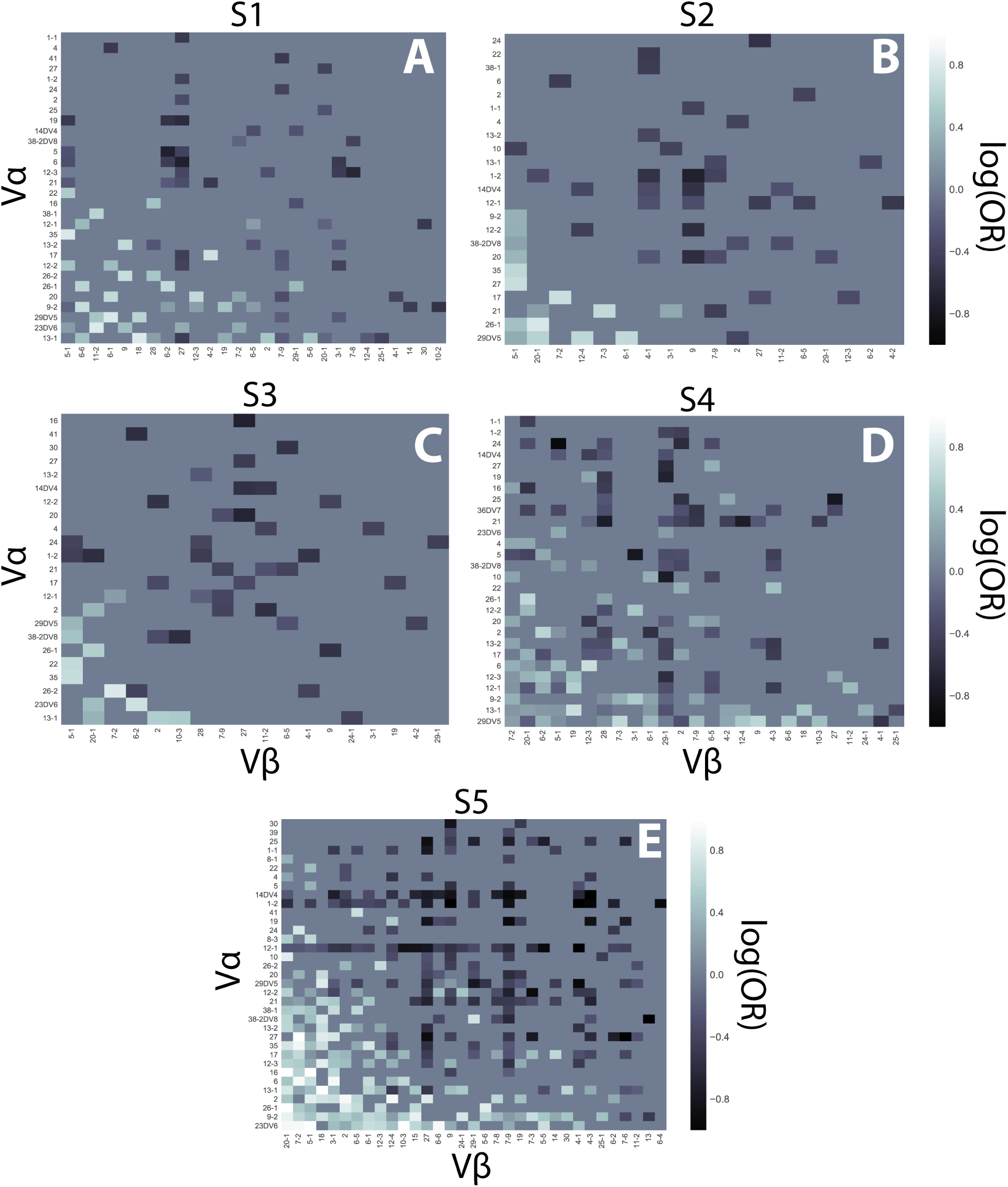
Paired V*αβ* CD4^+^:CD8^+^ log odds ratios for each individual. CD4^+^:CD8^+^ log odds ratio heatmaps for significant (*q<*0.05 by Fisher’s Exact test after correction for multiple-hypothesis testing) V*αβ* pairs for each individual. **(A)** S1, **(B)** S2, **(C)** S3, **(D)** S4, **(E)** S5. S6 is excluded due to low sample sizes.

**Supplemental Figure 11:**
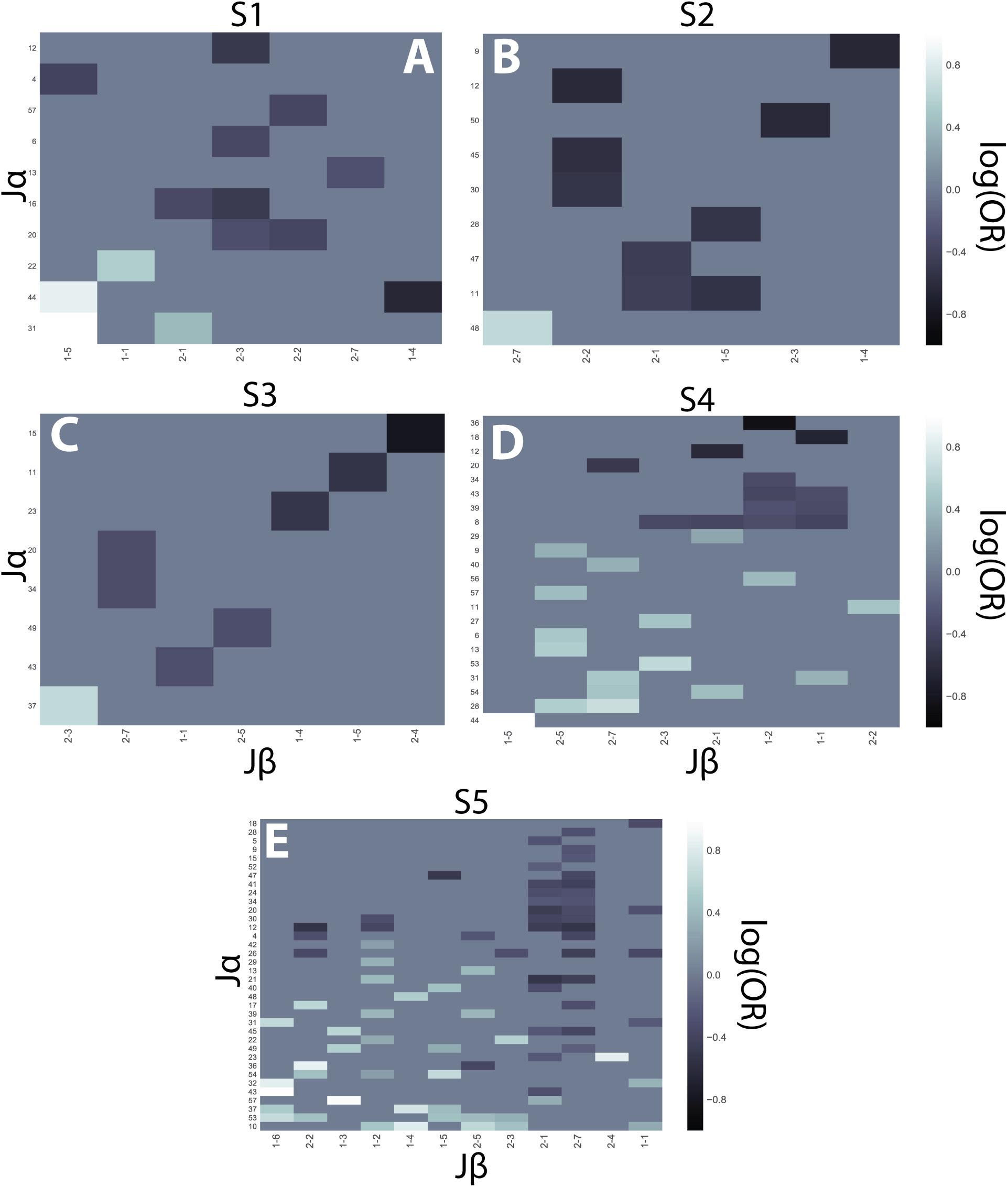
Paired J*αβ* CD4^+^:CD8^+^ log odds ratios for each individual. CD4^+^:CD8^+^ log odds ratio heatmaps for significant (*q<*0.05 by Fisher’s Exact test after correction for multiple-hypothesis testing) J*αβ* pairs for each individual. **(A)** S1, **(B)** S2, **(C)** S3, **(D)** S4, **(E)** S5. S6 is excluded due to low sample sizes.

**Supplemental Figure 12:**
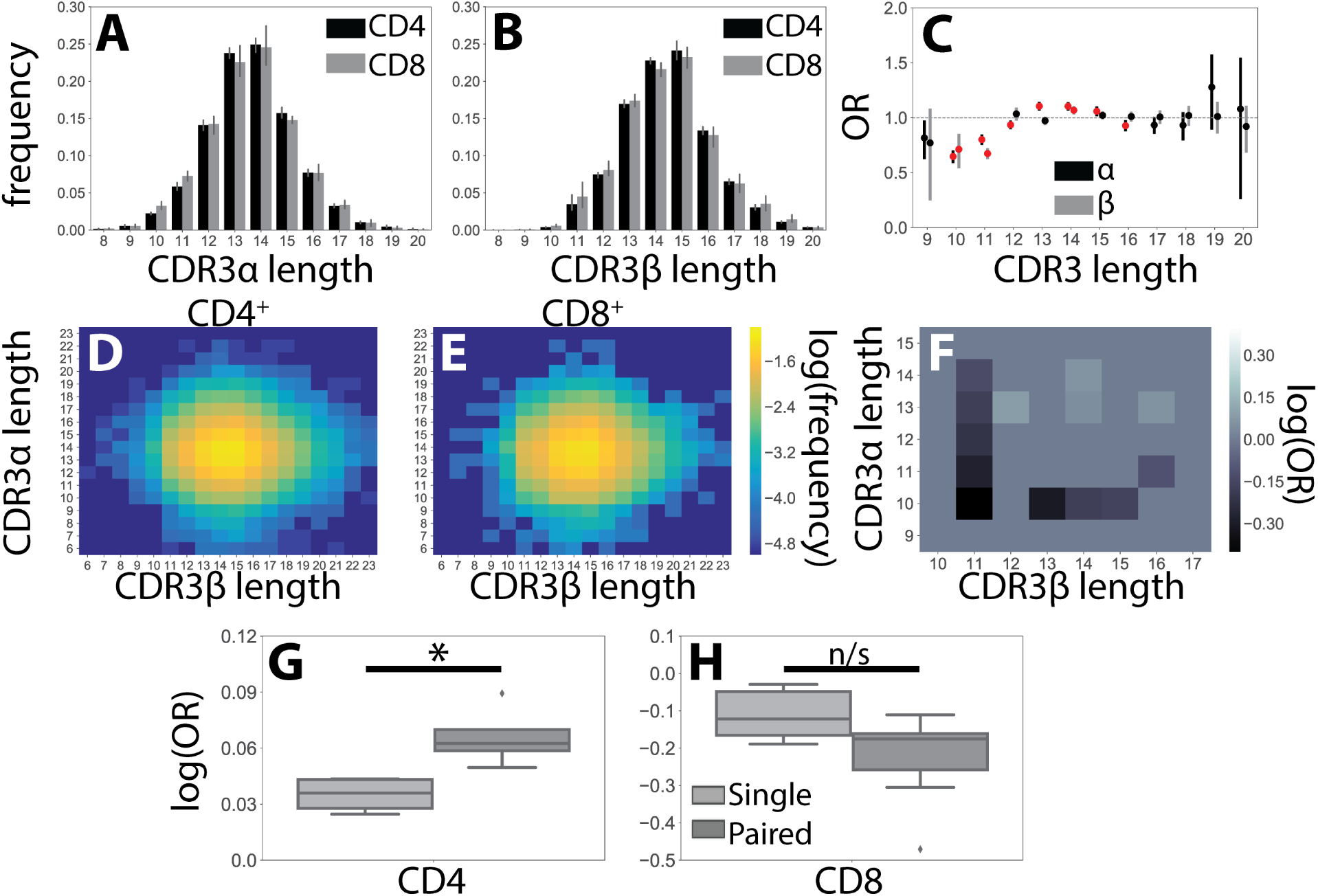
CDR3 length is weakly associated with T-cell lineage. CDR3 length histograms show no large differences between the CD4^+^ and CD8^+^ lineages for the **(A)** *α* and **(B)** *β* chain repertoires. Error bars represent standard deviation across individuals. **(C)** Odds ratios (OR) with 95% confidence intervals are shown for the *α* (gray) and *β* (black) single-chain repertoires. Red indicates statistical significance at the *p <* 0.05 level after Bonferroni correction.**(D)** Heatmaps showing log frequency of CDR3*αβ* length pairs within the CD4^+^ and **(E)** CD8^+^ TCR repertoires. **(F)** Significant ORs (*p<*0.05 by Fisher’s exact test after Bonferonni correction) for CDR3*αβ* length pairs reveal 14 *αβ* length pairs associated with a significant CD4^+^:CD8^+^ bias. **(G)** Boxplots compare statistically significant ORs for single chain and paired chain CDR3 lengths for pairs with CD4^+^ or **(H)** CD8^+^ bias. Significance between groups calculated by Mann-Whitney U test. \**p<*0.05. n/s-not significant.

**Supplemental Figure 13:**
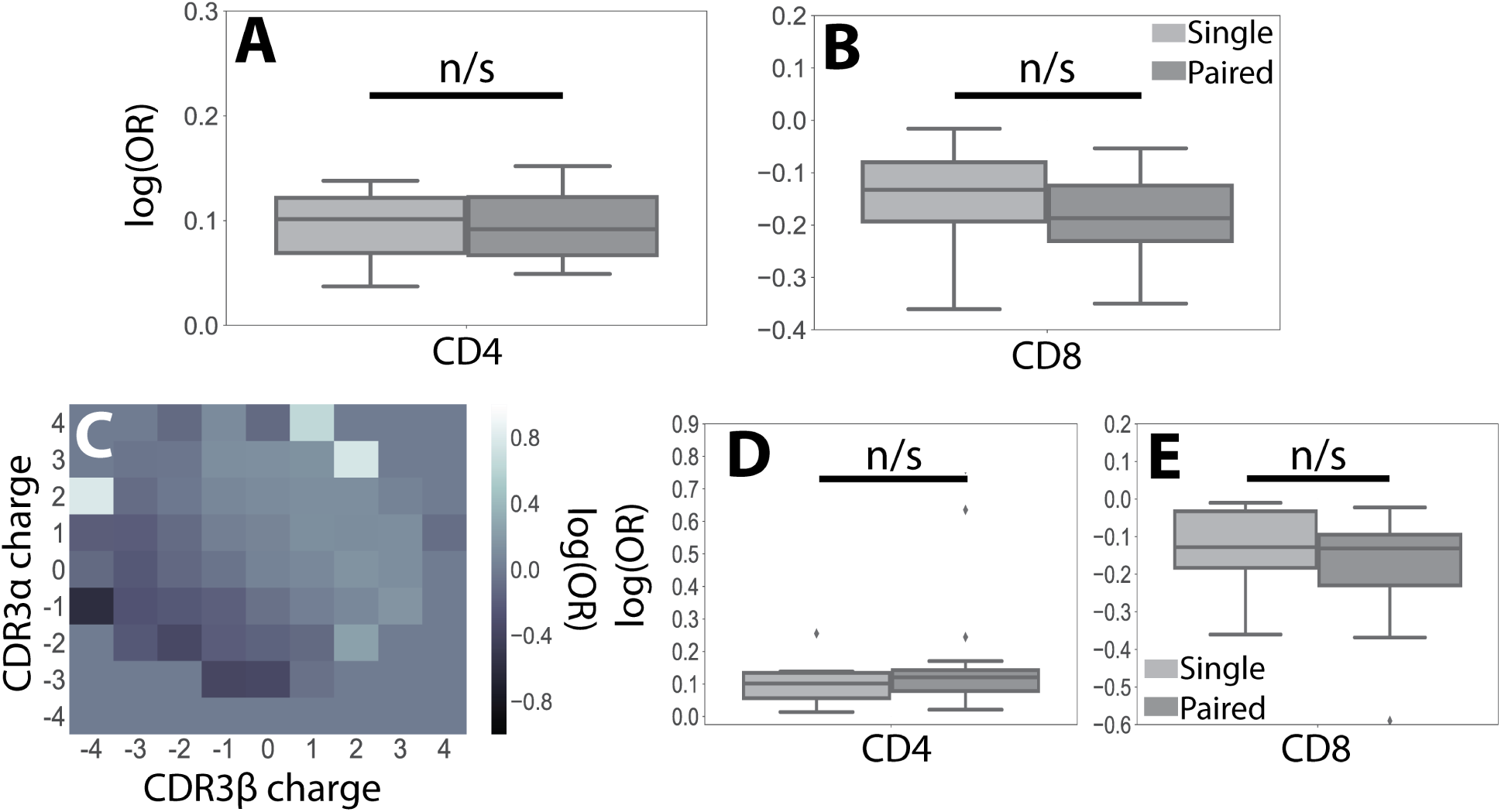
Paired CDR3 charges are more strongly associated with T-cell lineage than single chains. **(A)** All significant odds ratios associated with CDR3 charges for single chains (*α* or *β* alone) or for paired CDR3 charged (*αβ*) for CD4^+^ and **(B)** CD8^+^. **(C)** Heatmap for all paired CDR3*αβ* charges. **(D)** Boxplots representing the distribution of all CDR3*αβ* charge odds ratios with CD4^+^ and **(E)** CD8^+^ bias. n/s-not signficant.

**Supplemental Figure 14:**
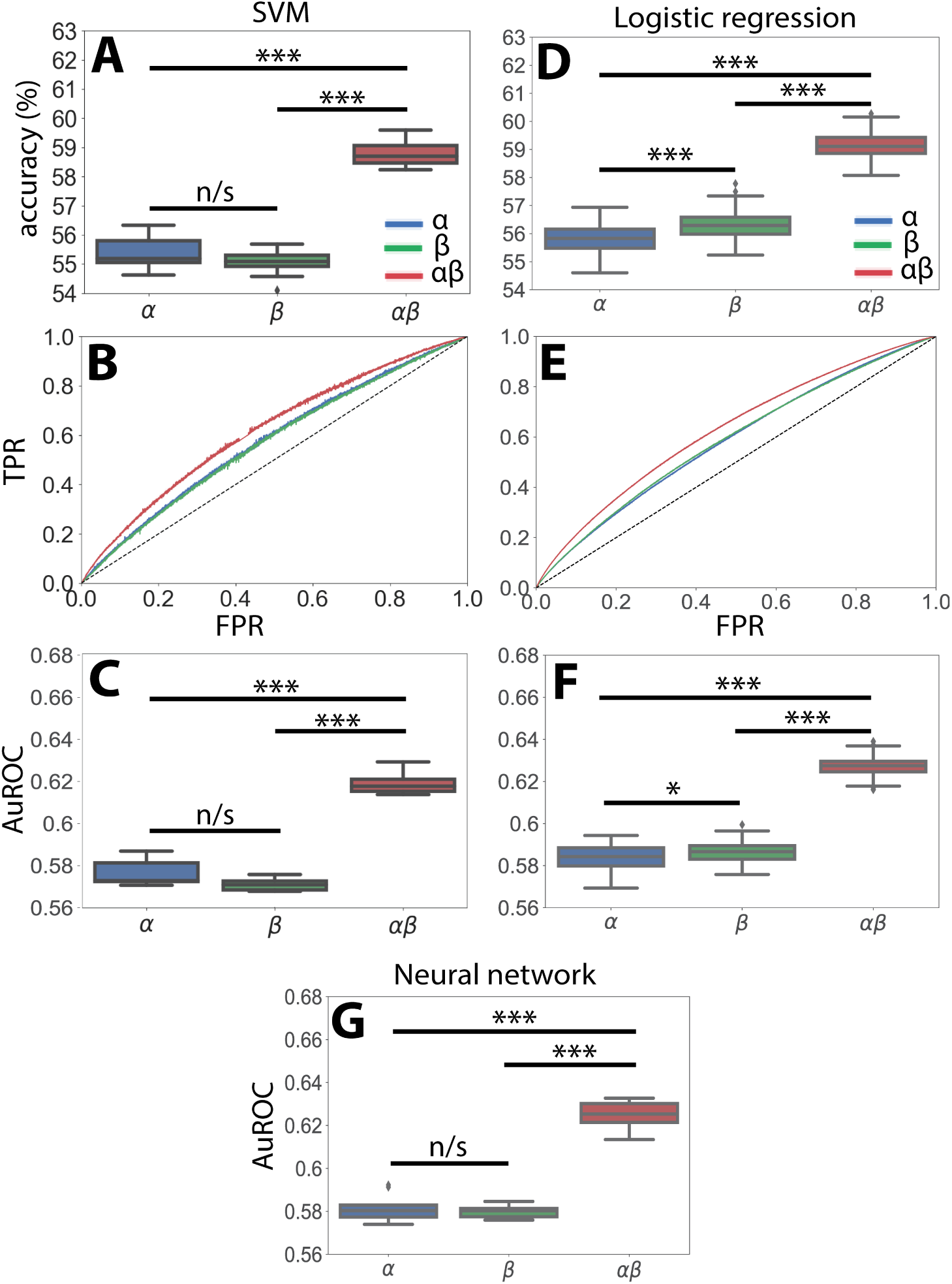
Accuracy of support-vector machine (SVM) and logistic regression models show *αβ* pairs outperform single chains. **(A)** SVM model accuracy for predicting T-cell CD4^+^ or CD8^+^ status from constant-length vectors encoding TCR features. **(B)** Receiver-operator curve (ROC) for SVM model shows *αβ* TCR pairs outperform either of the single chains alone. **(C)** Boxplots showing Area under the ROC (AuROC) for SVM classifier. **(D)** Similar results were obtained for logistic regression accuracy, **(E)** receiver-operator curve, and **(F)** AuROC. **(G)** AuROC for neural network trained in Figure 4.

**Supplemental Figure 15:**
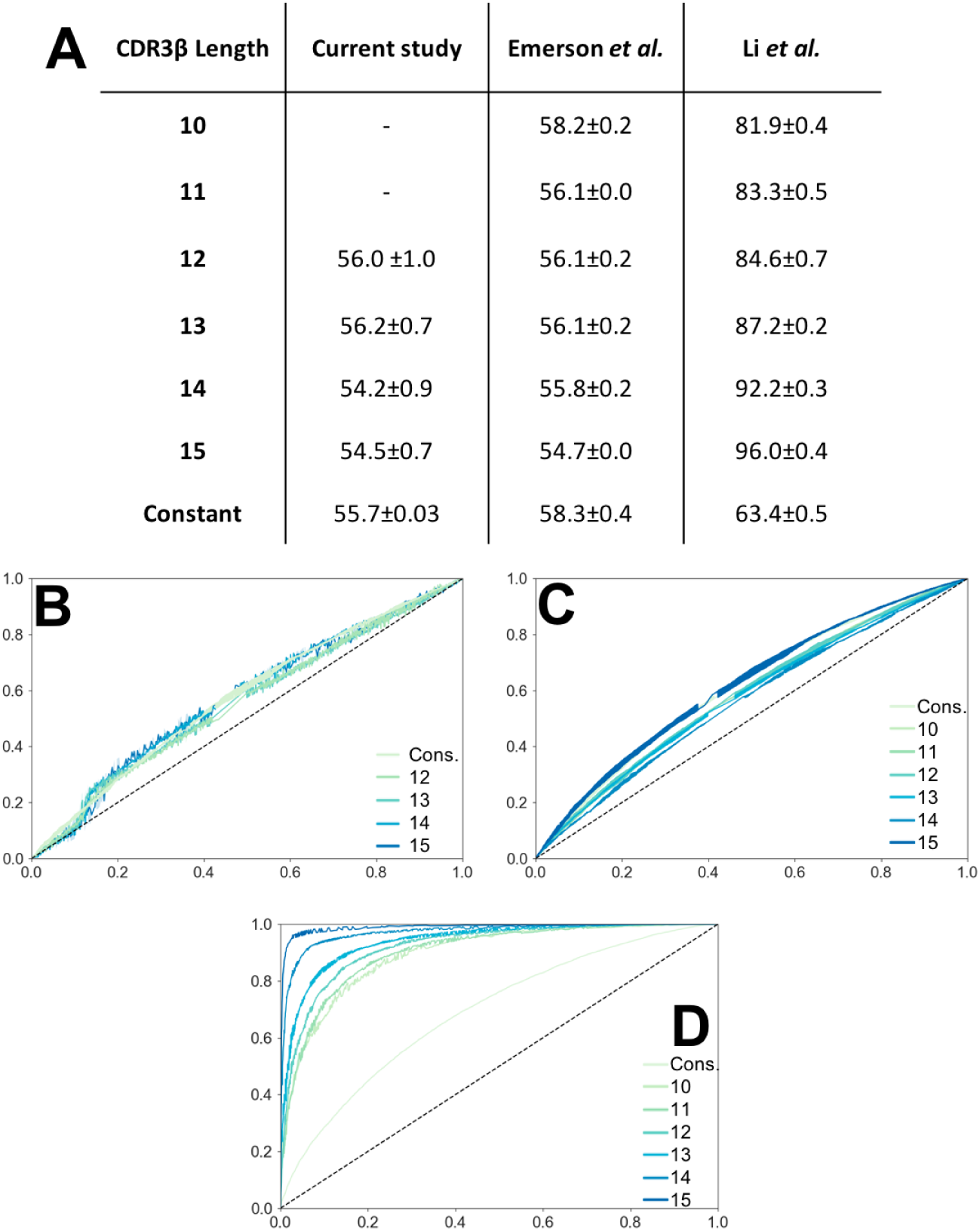
SVM trained on CDR3*β* sequences converted to Atchley factors. A support vector machine (SVM) was trained on vectors composed of CDR3*β* sequences converted into numerical array according to their Atchley factors. As these vectors are dependent on the length of the CDR3 sequence, SVMs were trained separately for CDR3 sequences of lengths between 10 and 15, as previously done^39^. For comparison, SVM accuracy for classifiers trained on CDR3*β* sequences converted to our constant length vector are also shown (Constant). **(A)** Accuracy for each model is reported as the percentage of correctly predicted CDR3 sequences using an independent testing set (25% of dataset). The Li *et al.* dataset is well described by this SVM model, with accuracy as high as 96%. However, this model fails to accurately describe either the dataset used in this study or that of Emerson *et al.* **(B)** Receiver operator curves (ROC) for the current dataset, **(C)** the Emerson *et al.* dataset, and **(D)** the Li *et al.* dataset show length-dependent SVMs accurately predict the Li *et al.* dataset, but fail to do so for the other two datasets.

**Supplemental Figure 16:**
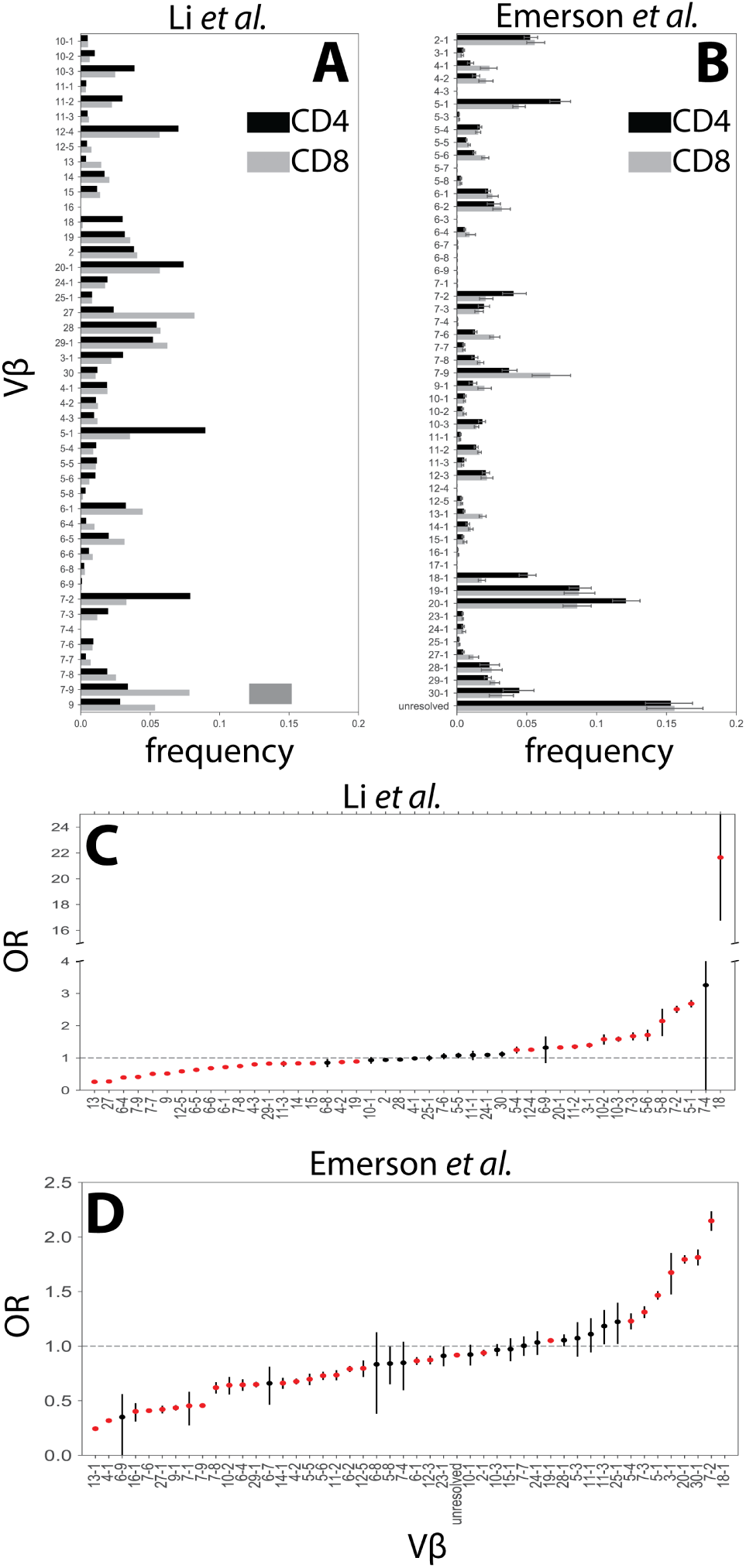
V region usage patterns vary substantially between the Li *et al.* and Emerson *et al.* datasets. **(A)** *β* TCR sequences were obtained from 621,085 CD4^+^ and 64,725 CD8^+^ cells previously by Li *et al.* ^39^. Comparison of V-usage frequencies for each germline region reveals large differences between the CD4^+^ and CD8^+^ repertoires in this dataset. **(B)** V-usage frequencies observed by comparing 3,212,682 CD4^+^ and 1,774,260 CD8^+^ TCR sequences taken from Emerson *et al.* reveal less variation between the two cell types ^40^. and more closely resemble the results obtained in the present study (Figure 2A-B). **(C)** We quantified the difference in V segment use in the CD4^+^ and CD8^+^ populations by calculating the odds ratio (OR) for each V region in the Li *et al.* dataset and **(D)** the Emerson *et al.* dataset independently. As observed from the frequency distributions, the Li *et al.* (OR: ~0.1-22.5)had substantially higher ORs associated with V region use as opposed to the Emerson *et al.* dataset.

**Supplemental Figure 17:**
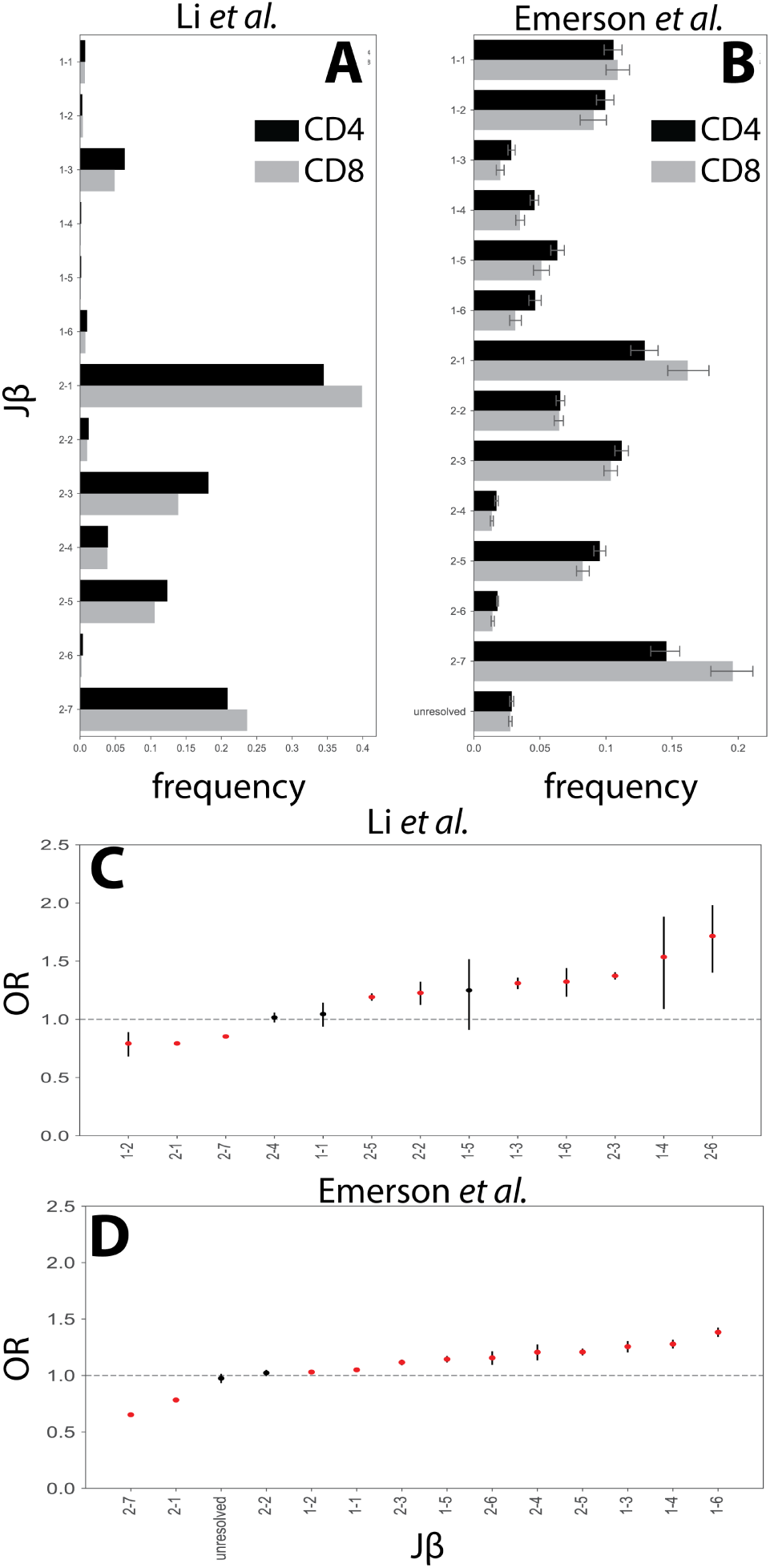
J region usage patterns vary substantially between the Li *et al.* and Emerson *et al.* datasets. **(A)** *β* TCR sequences were obtained from 621,085 CD4^+^ and 64,725 CD8^+^ cells previously by Li *et al.* ^39^. Comparison of J-usage frequencies for each germline region reveals large differences between the CD4^+^ and CD8^+^ repertoires in this dataset. **(B)** J-usage frequencies observed by comparing 3,212,682 CD4^+^ and 1,774,260 CD8^+^ TCR sequences taken from Emerson *et al.* reveal less variation between the two cell types ^40^. and again more closely resemble the results obtained in the present study (Sup. Fig. 4A-B) **(C)** We quantified the difference in J segment use in the CD4^+^ and CD8^+^ populations by calculating the odds ratio (OR) for each J region in the Li *et al.* dataset and **(D)** the Emerson *et al.* dataset independently. As observed from the frequency distributions, the Li *et al.* (OR: ~0.1-22.5) had substantially higher ORs associated with J region use as opposed to the Emerson *et al.* dataset.

**Supplemental Figure 18:**
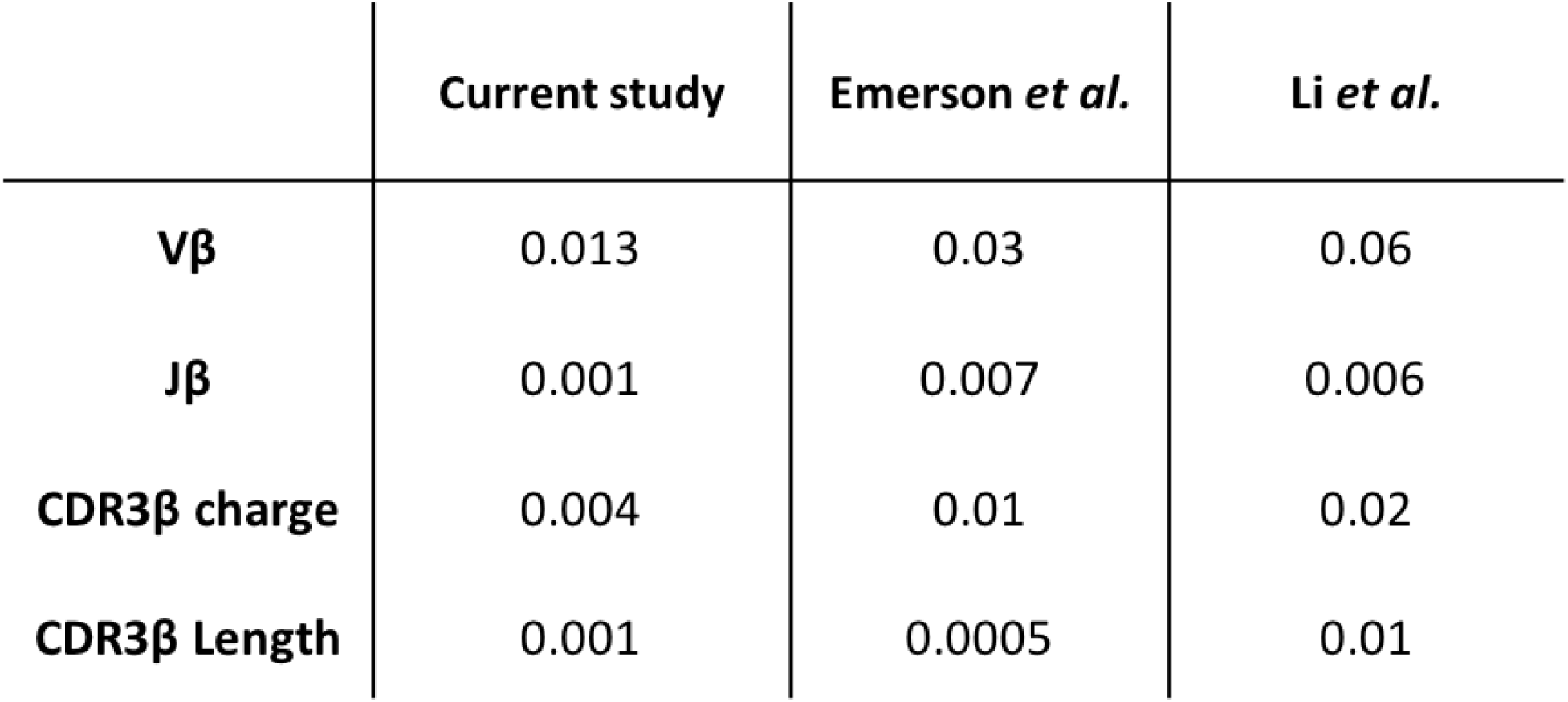
Mutual information between chain features and T-cell lineage for each dataset. Mutual information with finite sampling correction was calculated for the association between *β* chain features (V*β*, J*β*, CDR3*β* length and charge) and lineage for the dataset used in this study (from Table 1), by Emerson *et al.* and Li *et al.* ^39,40^ Substantially higher mutual information values, indicating stronger associations, were found for the Li *et al.* dataset as compared to the other two datasets.

